# The E3 ubiquitin ligase DTX3L and the deubiquitinase USP28 fine-tune DNA double strand repair through mutual regulation of their protein levels

**DOI:** 10.1101/2023.01.30.526213

**Authors:** Daniela Mennerich, Yashwanth Ashok, Carlos Vela-Rodríguez, Heli I. Hentilä, Melanie Rall-Scharpf, Lisa Wiesmüller, Renata Prunskaite-Hyyryläinen, Lari Lehtiö, Thomas Kietzmann

**Affiliations:** Faculty of Biochemistry and Molecular Medicine and Biocenter Oulu, University of Oulu, Oulu, Finland; Department of Obstetrics and Gynecology, Ulm University, Ulm, Germany

**Keywords:** Ubiquitination, DTX3L, USP28, hypoxia, c-MYC

## Abstract

The DNA damage response (DDR) relies on a complex protein network to maintain genomic integrity, yet the interplay between post-translational modifiers remains poorly understood. Here, we uncover a novel regulatory axis between the E3 ubiquitin ligase DTX3L and the deubiquitinase USP28 at DNA double-strand breaks (DSBs). Our results reveal a sophisticated feedback mechanism in which DTX3L ubiquitinates USP28, leading to its proteasomal degradation, while USP28 counteracts by deubiquitinating both itself and DTX3L. This cross-regulation fine-tunes DSB repair in multiple pathways, including non-homologous end joining (NHEJ), homologous recombination (HR), single-strand annealing (SSA), and microhomology-mediated end joining (MMEJ). Strikingly, the detrimental effects of USP28 depletion on these repair pathways were rescued by concurrent DTX3L knockdown. Collectively, our work uncovers a novel layer of DDR regulation in which DTX3L and USP28’s antagonistic activities calibrate cellular responses to genotoxic stress, thus identifying promising therapeutic targets to combat diseases associated with genomic instability.

**Highlights:**

- DTX3L and USP28 physically interact and colocalize in cellular sub-compartments, with the N-terminal D1-D3 domains of DTX3L primarily mediating the interaction
- DTX3L ubiquitinates USP28 for degradation, while USP28 deubiquitinates itself and DTX3L, creating a sophisticated feedback mechanism.
- The DTX3L-USP28 circuit influences levels of key proteins like HIF-1α, p53, and c-MYC, suggesting broader impacts on cellular stress responses.
- DTX3L and USP28 cooperatively regulate multiple DSB repair pathways, including NHEJ, HR, SSA, and MMEJ, with USP28 depletion effects rescued by DTX3L silencing.

## Introduction

Ubiquitination is a posttranslational protein modification that involves the covalent attachment of ubiquitin molecules to target proteins, leading either to their degradation by the proteasome or the regulation of a vast array of cellular processes.

Three different types of ubiquitinations are known: (a) monoubiquitination, (b) multi-monoubiquitination (c) polyubiquitination. While monoubiquitination mainly affects protein localization, polyubiquitination is best known for its regulation of protein abundance by promoting proteasomal degradation of polyubiquitinated proteins. However, it has recently emerged that polyubiquitination can have non-proteolytic functions by taking part in transcription regulation, inflammation, endocytosis, mitophagy, cell division and DNA repair ^1^. While the sequential action of three enzymes (E1, E2, and E3) is required for successful attachment of ubiquitin to a substrate, the substrate specificity is lastly determined by the E3 ubiquitin ligase ^1^.

Apart from the actual ubiquitination, there are other ubiquitin-like modifiers (UBLs) such as NEDD8, SUMO, and ISG15 that can also be conjugated to ubiquitin and target proteins to modulate their activity and localization. The latter may lead to formation of hybrid chains consisting, for example, of both ubiquitin and NEDD8 molecules. Overall, the presence of multiple types of UBLs and the formation of hybrid chains adds complexity to the regulation of protein degradation and cellular signaling ^2^.

Recently, one member of the DTX family of E3 ubiquitin ligases ^3^, DTX3L, has emerged to be involved in several diseases such as inflammation and cancer that are linked to differentiation, apoptosis, and DNA repair. However, the knowledge of DTX substrates and their molecular mechanisms is limited. DTX3L was originally identified as a binding partner of PARP9 (BAL1/ARTD9) and linked to DNA repair due to its ability to monoubiquitinate histone 4 and to promote recruitment of the tumor suppressor (TP53BP1) to DNA damage sites ^4^. In addition to DTX3L, there are other human E3 ligases (DTX1, DTX2, DTX3, and DTX4) which share a common C-terminal DTC domain that^5^ mediates the formation of a covalent linkage between the C-terminus of ubiquitin and the adenosine-proximal ribose of ADPr, generating unique Ub-ADPr hybrid chains ^6,7^. Moreover, DTX3L was found to be able to undergo autoubiquitination and to carry out poly-ubiquitination with different ubiquitin linkages ^8,9^. However, the functional relevance especially with DNA repair remains to be determined ^10^.

Deubiquitinases (DUBs) comprise a vast group of proteases that cleave ubiquitin from substrates or process ubiquitin precursors to generate free ubiquitin pools ^11^. In humans, there are about 100 enzymes that are grouped into seven main classes based on their sequence relationship and structural fold. These distinct classes of DUBs highlight the diversity and complexity of the deubiquitination process in cellular regulation ^12,13^.

Although some general understanding about the function of DUBs exists, not much is known about the specific roles of many family members. From those, ubiquitin-specific protease 28 (USP28) encoded at chromosome 11q23 has recently gained interest as it was implicated to modulate several cellular processes such as DNA damage response, cell-cycle, and tumorigenesis. So far, two USP28 isoforms could be identified; the ubiquitously expressed canonical isoform and the 62 amino acids longer tissue-specific isoform that is found in muscles, heart and brain ^14^.

Like with DTX3L, the biological role of USP28 is connected to the DNA damage response, which is illustrated by the fact that USP28 is recruited to DNA damage sites ^15,16^. USP28‘s activity in response to DNA damage is tightly regulated by the ATM kinase that phosphorylates serine 67 and 714 leading to an increase in its catalytic activity ^16^. By contrast, cleavage of USP28 by caspase-8 causes loss of its catalytic activity and progression of the cell-cycle beyond the p53-dependent G2/M DNA damage checkpoint ^17^. Similarly, sumoylation of lysine 99 in the N-terminus of USP28 was shown to reduce USP28 activity, whereas the SUMO protease SENP1 could reverse this effect under hypoxic conditions ^18^. In addition, USP28 was shown to interact with TP53BP1 and to rescue it from degradation along with other proteins involved in the DNA damage response such as p53, claspin, and CHK2 ^16,19,20^. Within this scenario, the specific recruitment of the USP28 interactor TP53BP1 to DNA damage sites, specifically to DNA double strand breaks (DSBs) is promoted by a heterodimeric complex consisting of the ADP-ribosyltransferase PARP9 and the E3 ubiquitin ligase DTX3L ^9^. Considering this, we hypothesized that USP28 could oppose the action of DTX3L; Vice versa DTX3L could potentially ubiquitinate and affect USP28 function or levels. Consequently, this interplay could impact DNA DSB repair.

Here we uncover that USP28 is indeed able to deubiquitinate DTX3L. At the same time, we show that DTX3L ubiquitinates USP28 and promotes its proteasomal degradation. We also show that regulation of USP28 by DTX3L has functional consequences on known USP28 substrates such as TP53, c-MYC and HIF-1α as well as on DSB repair pathway usage. Together, these data indicate that the interplay between USP28 and DTX3L is critical for the DNA damage response and its perturbation during tumorigenesis.

## Results

### DTX3L and USP28 show physical interaction

To expand our understanding of the relationship between DTX3L and USP28, we first performed immunofluorescence microscopy experiments to test whether the two proteins colocalize. The analyses show that a small fraction of both proteins localize to the cytoplasm whereas the majority was found in the nucleus and as indicated by a focal pattern particularly to certain sub-compartments (**Fig. S1A**). Given that DTX3L has been previously implicated in recruiting TP53BP1 ^4^, we hypothesized that DTX3L, USP28 and TP53BP1 are components of the same complex. To test this hypothesis, we performed immunoprecipitation experiments using control IgG or antibodies against TP53BP1 and USP28 in SK-MES-1 lung cancer cells. The results showed that both DTX3L and USP28 were present in the TP53BP1 immunoprecipitates **(Fig. S1B)**.

To further investigate the potential direct interaction between DTX3L and USP28, we next employed a proximity ligation immunoassay (PLA) and co-immunoprecipitation analyses. While the co-immunoprecipitation analyses from whole cell lysates verify the interaction of endogenous DTX3L and USP28, the PLA assay revealed fluorescence signals indicating an interaction between USP28 and DTX3L in both, the cytoplasm and in the nucleus (**Fig. 1A,B,C**). To provide more robust evidence for the interaction and colocalization of USP28 and DTX3L in both compartments, we performed additional co-immunoprecipitation from cytoplasmic and nuclear fractions. The data show that the two proteins can localize and interact in both compartments, *i.e.*, the cytoplasm and the nucleus (**Fig. 1D**).

**Figure 1.**
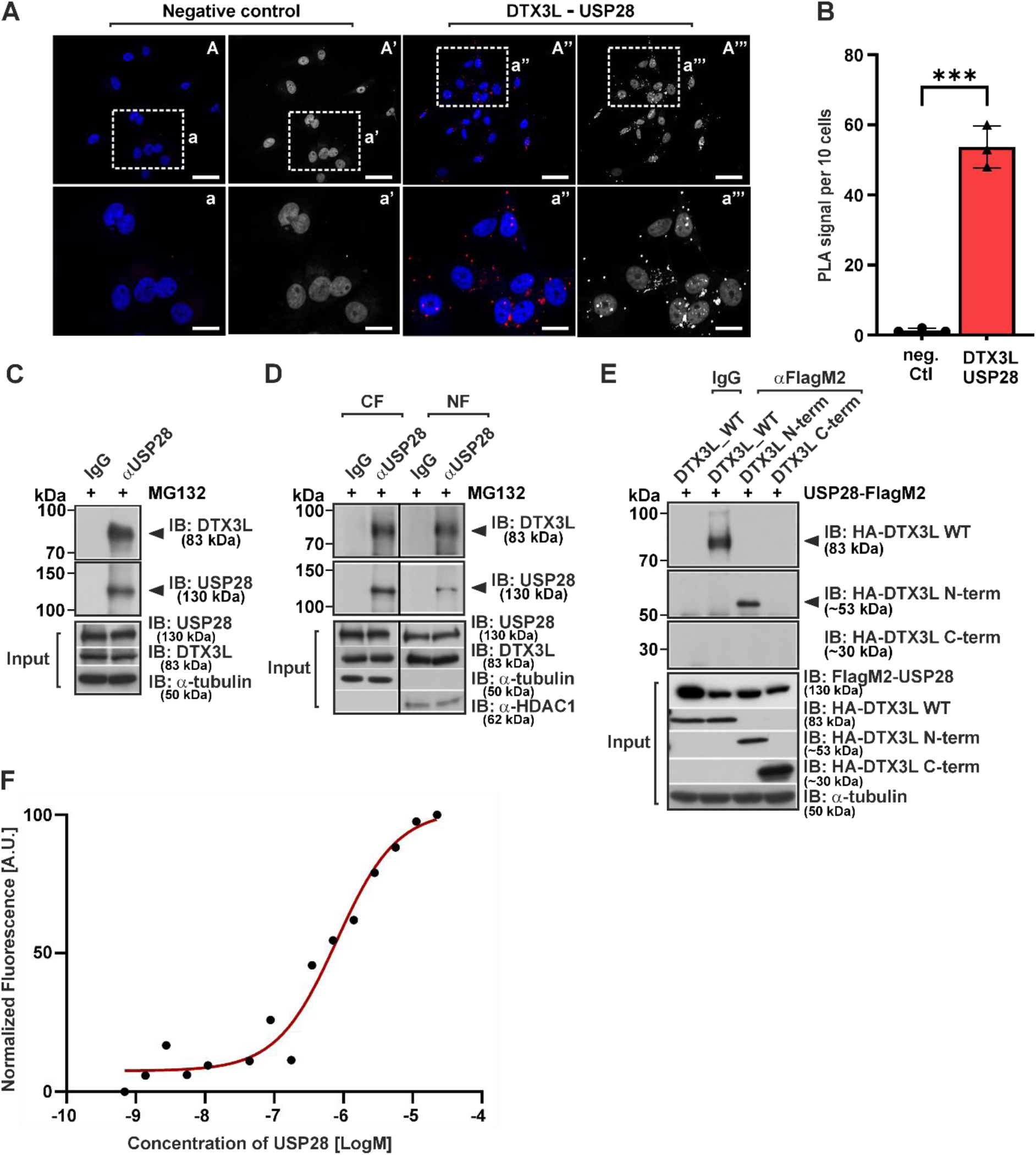
DTX3L and USP28 interact with each other. **(A)** Interaction of endogenous DTX3L and USP28 was visualized by proximity ligation assay (PLA) in SK-MES-1 cells. Cells were grown on cover slips, fixed and immunostained according to Duolink manufacturés protocol. Red spots reflect the interaction. DAPI stained nuclei are shown in blue. Scale bars: A-A’’’ 50 μm and a-a’’’ 20 μm **(B)** Quantification of the number of PLA signals. ****significant difference between the number of proximity ligation sites in the negative control vs DTX3L/USP28. Data are mean +/-SD from 3 independent experiments. The statistical significance of differences was determined by ordinary one-way ANOVA. ***p<0.001. **(C,D)** Cell extracts from PC-3 cells were immunoprecipitated with either IgG control or USP28 antibodies. Resulting precipitates were first analyzed by immunoblotting (IB) using a DTX3L; thereafter, the membrane was re-probed with a USP28 antibody. Whole cell lysate (C) and cytoplasmic (CF) and nuclear fraction (NF) (D). **(E)** HEK293 cells were cotransfected with expression vectors encoding full length Flag-tagged USP28 and full-length HA-tagged DTX3L or the N-terminal HA-tagged DTX3L (A2-A516) deletion mutant or the C-terminal HA-tagged DTX3L (K555-E740) deletion mutant. Cell extracts were immunoprecipitated with either IgG control or USP28 antibodies. Immunoprecipitates were analyzed by immunoblotting (IB) using a HA-tag antibody. **(F)** Representative MST binding curve for DTX3L and USP28 interaction. All binding curves are shown in Fig. S3.

We also performed co-immunoprecipitations by using HEK293 cells that were transfected with vectors allowing expression of HA-tagged DTX3L and Flag-tagged USP28. After USP28 was immunoprecipitated with a FlagM2-tag antibody, the subsequent western blot analysis with HA-tag antibodies revealed presence of DTX3L in the precipitates (**Fig. S1C**).

Next, we conducted additional co-immunoprecipitations to gain insight into the interaction interfaces of the two proteins. We used cells expressing either the HA-tagged DTX3L D1-D3 domains (A2-A516), i.e., the N-terminal part, or the RING-DTC domain (K555-E740) C-terminal part of DTX3L. Interestingly, our data revealed that the interaction primarily occurs between USP28 and the N-terminal part of DTX3L, specifically involving the D1-D3 domains (A2-A516), rather than the RING-DTC domains (K555-E740) (**Fig. 1E**).

To further investigate the direct USP28-DTX3L interaction and to determine their binding affinity in solution, we next expressed both proteins recombinantly and performed interaction measurements with microscale thermophoresis (MST). Therein, we titrated fluorescently labelled DTX3L with USP28 to determine their dissociation constant (K_d_). CD spectra for USP28 were recorded to assess the degree of folding of the recombinantly produced proteins (**Fig. S2**). Although not all MST curves reached complete saturation due to the high protein concentration required, the data show that the K_d_ value was 4.9 µM (LogK_d_ = 5.5 ± 0.5) (**Fig. 1F, Fig. S3**). MST trace reading at different USP28 concentrations showed that there would not be non-specific binding interfering with the measurement **(Fig. S4)**. Together, the data support the view that USP28 and DTX3L can undergo complex formation.

### USP28 can remove poly-ubiquitin (poly-Ub) chains catalyzed by DTX3L

Having shown that DTX3L and USP28 can even reside in the same complex and knowing that DTX3L undergoes auto-ubiquitination from our previous report ^8^ we sought to investigate whether USP28 could catalyze the removal of poly-Ub chains from DTX3L. To test this idea, we used full-length DTX3L as a substrate in an *in vitro* ubiquitination assay. In agreement with our previous report, the assay revealed that DTX3L was able to undergo auto-ubiquitination (**Fig. 2A-C lane 8, Fig. S5A-B lane 7**). In the next experiments, we increased the reaction time for DTX3L and added USP28 later while keeping the total reaction time constant. The data reveals that addition of wild-type USP28 to the enzyme assay was indeed able to reverse the auto-modification of DTX3L (**Fig. 2A-C lane 9, Fig. S5A-B lane 9, 11, 13**). Furthermore, when USP28 was added to the reaction at different concentrations and different times, deubiquitination of DTX3L occurred in a time- and concentration-dependent manner **(Fig.S5C,D).** In contrast, USP28^C171A^, a catalytically inactive mutant, was unable to do so (**Fig. 2A-C lane 10**). The deubiquitination activity of USP28 did neither require the N-terminal region consisting of the ubiquitin-binding domains (UBA), the ubiquitin-interaction motif (UIM) and the SUMO-interaction motifs (SIM), nor the C-terminal extension as a fragment encompassing only the catalytic domain (USP28^cat^, E147-L652) was sufficient to remove the ubiquitins from DTX3L (**Fig. 2A-C lane 11**). Thus, these results show that USP28 can completely remove the ubiquitin modifications from DTX3L *in vitro*.

**Figure 2.**
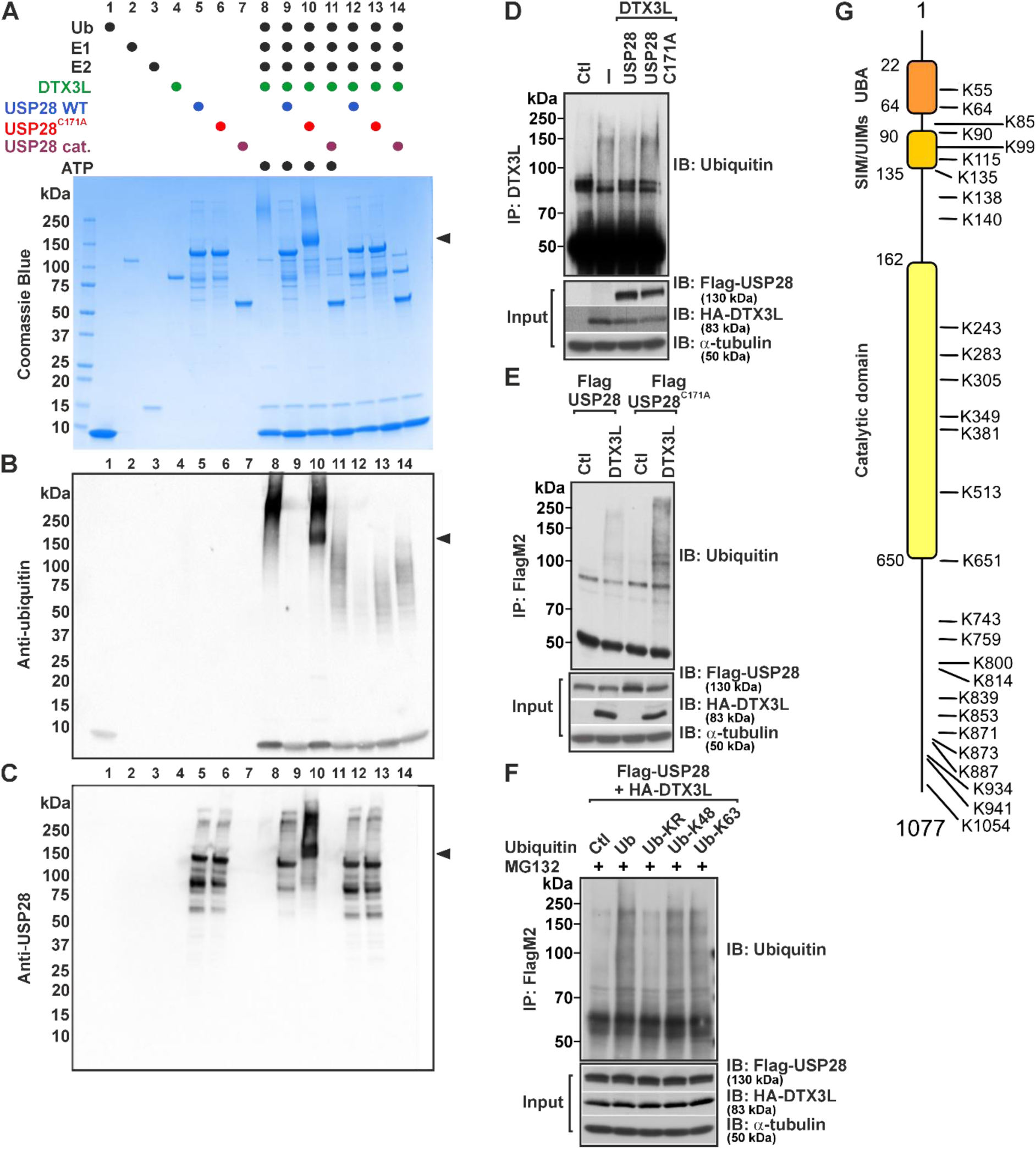
Functional interplay between USP28 and DTX3L. **(A)** *In vitro* ubiquitination assay indicating the auto-ubiquitination activity of DTX3L and deubiquitinating activity of USP28. Protein concentrations used were Ub (50 µM), E1 (0.4 µM), UbcH5a (E2) (2 µM) and DTX3L (0.7 µM). USP28 was used at a final concentration of 3 µM. SDS-PAGE showing appearance of the DTXL3 ubiquitination pattern that is visible as a smear (lane 8) and its disappearance due to hydrolysis by USP28 (lane 9) and/or the USP28 catalytic domain (lane 11). Ubiquitinated DTX3L appears as a high molecular weight smear and the black arrow indicates a band that corresponds to ubiquitinated USP28. Note, that for unknown reasons USP28 preparations showed some degradation (lanes 5-6). **(B)** Western blot of the assay in panel A probed with an anti-ubiquitin antibody. The black arrow indicates a band that corresponds to ubiquitinated USP28. **(C)** Western blot of the assay in panel A probed with anti-USP28. The arrow indicates a band shift of the ubiquitinated catalytically inactive mutant USP28^C171A^ in the presence of ATP, Ubiquitin (Ub), E1 and E2 ligases as well as DTX3L. **(D, E, F)** Cell based ubiquitination assays. HEK293 cells were transfected with an empty vector (Ctl) or expression vectors for DTX3L, wild type USP28 and the catalytically inactive mutant USP28^C171A^. In D, ubiquitinated DTX3L was detected with ubiquitin antibody after immunoprecipitation of DTX3L. In E, ubiquitinated USP28 was detected with ubiquitin antibody after immunoprecipitation of USP28 with a FlagM2-tag antibody. In F, cells were transfected with DTX3L and Flag-USP28 expression vectors as well as with empty vector (Ctl) or vectors for wild type ubiquitin (Ub), a mutant in which all lysine residues were converted to arginine (Ub-KR) or mutants where either only K48 (Ub-K48) or K63 (Ub-K63) remained intact. Ubiquitinated USP28 was detected with ubiquitin antibody after immunoprecipitation of USP28 with a FlagM2-tag antibody. **(G)** Domain organization of USP28 and the ubiquitination sites identified by mass spectrometry using the USP28^C171A^ mutant. UBA, - ubiquitin-associated domain; UIM, -ubiquitin interaction motif; SIM,-SUMO interaction motif.

We next aimed to corroborate the results from the *in vitro* assay in cells. To this end, we performed DTX3L ubiquitination assays in HEK293 cells upon overexpression of DTX3L along with wild-type USP28 and the catalytically inactive USP28^C171A^ mutant. After DTX3L immunoprecipitation and western blot analysis with a ubiquitin antibody it was evident that overexpression of DTX3L alone increased the appearance of the polyubiquitin smear when compared to empty vector (Ctl) transfected cells (**Fig. 2D**); a finding in agreement with the autoubiquination of DTX3L in the in vitro assays. Furthermore, we found that wild type USP28 was able to reduce the appearance of the ubiquitin chains from DTX3L whereas the inactive mutant USP28^C171A^ could not (**Fig. 2E**). Together, these data indicate that USP28 can act as a DUB on DTX3L.

### DTX3L ubiquitinates USP28

In addition to the auto-ubiquitination of DTX3L, we also observed that DTX3L could ubiquitinate the USP28^C171A^ mutant (**Fig. 2 lane 10**). This was only seen with the inactive USP28^C171A^ mutant (**Fig. 2A, 2B lane 10)** because the active USP28 would deubiquitinate i**A, 2B**tself (**Fig. 2A, 2B lane 12**) as well as DTX3L (**Fig. 2A, 2B lane 9**). When membranes were probed with an anti-ubiquitin antibody, the modified USP28^C171A^ became visible as a band that migrates faster in comparison to auto-ubiquitinated DTX3L and represents modified USP28^C171A^ (**Fig. 2B lane 10, black arrow**). To further confirm this, we probed the membrane with an anti-USP28 antibody. As expected, the blots reveal a shift of the band representing USP28^C171A^ towards a higher molecular weight as result of the ubiquitination (**Fig. 2C lane 10 versus lane 6**). Similarly, when we performed ubiquitination assays for USP28 in cells that either overexpressed wild type USP28 or the inactive USP28^C171A^ along with DTX3L, we found that DTX3L promoted ubiquitination of USP28. In line with the *in vitro* ubiquitination assay, this was especially evident in the cells expressing USP28^C171A^ (**Fig. 2E**). To further investigate the role of DTX3L’s E3 ligase activity in the regulation of USP28, we conducted additional experiments using an E3 ligase-deficient mutant of DTX3L. We used this mutant in both cell assays and with recombinant proteins and demonstrated that the inactive DTX3L does not ubiquitinate USP28 **(Fig. S6)**. Importantly, the cell-based data also indicate that this lack of ubiquitination is not compensated by other E3 ligases.

To further investigate whether the ubiquitination of USP28 by DTX3L renders it susceptible for proteasomal degradation, we performed an additional ubiquitination assay in the presence of either wild type ubiquitin (Ub) or three specific ubiquitin mutants. In the first mutant, all lysine residues were converted to arginine (Ub-KR), preventing ubiquitination and, in particular, polyubiquitin chain formation. As expected, use of this mutant resulted in a loss of USP28 ubiquitination by DTX3L (**Fig. 2F**). In the second mutant (Ub-K48), only one lysine, *i.e.,* K48, remained intact. Therefore, only K48-linked polyubiquitin chains can be formed when this mutant is used. The assay shows that DTX3L is able to ubiquitinate USP28 in the presence of this mutant. In the third mutant (Ub-K63), only K63 remains intact allowing only K63-linked polyubiquitin chains, and DTX3L was also able to ubiquitinate USP28 with this mutant (**Fig. 2F**).

Next, we used mass spectrometry to identify the sites within USP28^C171A^ that become ubiquitinated in the presence of DTX3L. In total, 28 lysins were found to be modified by DTX3L and those ubiquitination sites were distributed over the whole protein without any domain preference (**Fig. 2G**). Together, the data show that USP28 can act as a DUB on DTX3L and vice versa, that USP28 is a substrate for DTX3L.

### DTX3L and USP28 mutually control their levels in cells

DTX3L is known to function as an E3 ubiquitin ligase, promoting the ubiquitination and subsequent degradation of interacting target proteins such as USP28. On the other hand, as a deubiquitinase USP28 can remove ubiquitin modifications leading to their stabilization. As the in vitro studies have shown that both DTX3L and USP28, appear to control each other’s ubiquitination, we next aimed to investigate whether this mutual interplay affects their stability in a broader physiological context in cells.

First, we wanted to test if DTX3L would regulate USP28 levels and vice versa whether USP28 would regulate DTX3L levels. To this end, we first overexpressed DTX3L in SK-MES-1 lung tumor cells as well as in HeLa cells and analyzed USP28 levels. As USP28 plays a role in various stress pathways including hypoxia signaling ^21^, we also exposed those cells to normoxia and hypoxia to examine changes in downstream USP28 substrates such as HIF-1α, p53, and c-MYC ^22^. When analyzing the cells, we observed that overexpression of DTX3L decreased USP28 levels under both normoxia and hypoxia (**Fig. 3A,B**) which is in agreement with polyubiquitination of USP28 by DTX3L. In line with previous observations the decrease of USP28 was accompanied by a reduction in HIF-1α ^21^, p53 ^23^, and c-MYC levels ^16,19,23,24^ ^25^. In addition, we performed knockdown experiments with two different DTX3L shRNAs. We found that knockdown of DTX3L raised USP28 protein levels up to two-fold and induced HIF- 1α levels under hypoxia, and p53, and c-MYC levels under both normoxia and hypoxia in SK-MES-1 cells (**Fig. 3A,B**) and in HeLa cells (**Fig. S7A**). Importantly, knockdown of DTX3L reduced its target levels without affecting USP28 mRNA levels (**Fig. S7B**).

**Figure 3.**
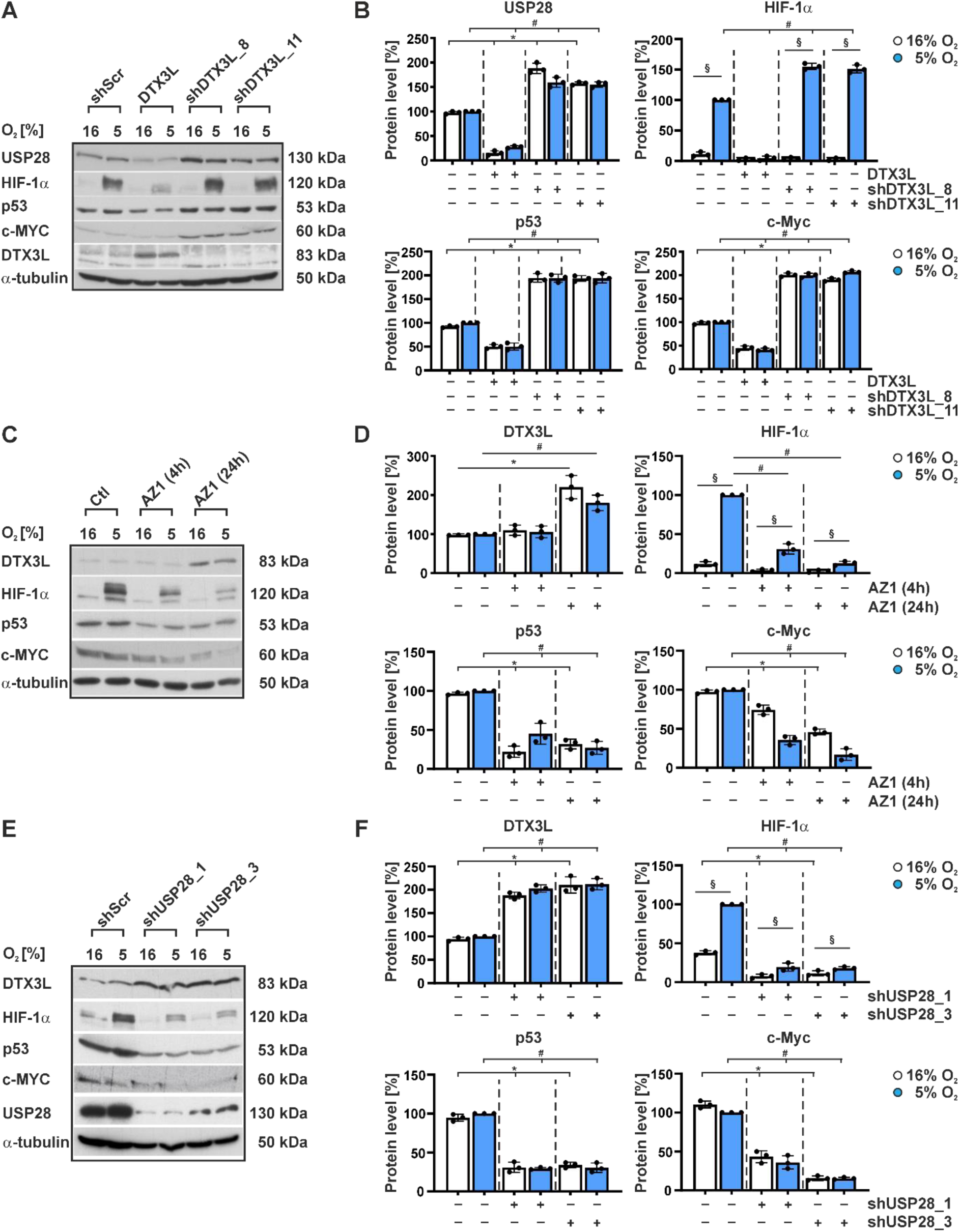
Mutual regulation of DTX3L and USP28 protein levels. **(A)** SK-MES-1 cells were transfected either with expression vector for scrambled control shRNA (shScr), or with expression vectors for full-length DTX3L or one of two independent shRNAs (shDTX3L_8, shDTX3L_11) against DTX3L. After transfection, cells were further cultured under normoxia (16% O_2_) or hypoxia (5% O_2_) for 4 hours. USP28, HIF-1α, p53, c-MYC, and DTX3L protein levels were measured by Western blot analysis. Alpha tubulin served as a loading control. **(C)** SK-MES-1 cells were treated with the USP28/25 inhibitor AZ1 (10 µM) and cultured under normoxia and hypoxia for 4 h and 24 h. DTX3L, HIF-1α, p53, and c-MYC protein levels were measured by Western blot analysis. **(E)** SK-MES-1 cells were transfected with expression vectors for scrambled control shRNA (shScr) or shRNA 1 or shRNA 3 against USP28. After transfection, cells were further cultured under normoxia or hypoxia for 4 h. DTX3L, HIF-1α, p53, c-Myc and USP28 protein levels were measured by Western blot analysis. **(B,D,F)** Quantification of USP28, DTX3L, HIF-1α, p53 and c-MYC. In each experiment the protein levels at 5% O_2_ control (Ctl) or shScr were set to 100%. *significant difference for 16% O_2_: Ctl vs DTX3L, vs shDTX3L, vs shUSP28 or vs AZ1, ^§^significant difference between 16% O_2_ vs 5% O_2_, **^#^**significant difference for 5% O_2_: Ctl vs DTX3L, vs shDTX3L, vs shUSP28 or vs AZ1, p < 0.05.

Next, we were interested to see the impact of USP28 on DTX3L. As USP28 can remove the auto-ubiquitination from DTX3L, it would at first glance be expected that inhibition or lack of USP28 would reduce DTX3L levels due to enhanced degradation. We tested this assumption by using two approaches. Firstly, we exposed cells to the dual selective USP28/25 inhibitor AZ1 ^26^ and secondly, we used two different shRNAs targeting USP28 ^21^. In contrast to our hypothesis, we found that both chemical as well as genetic inhibition of USP28 upregulated the levels of DTX3L whereas HIF-1α, p53, and c-MYC were downregulated in SK-MES-1 cells (**Fig. 3C-F**) and in HeLa cells (**Fig. S7C**). Furthermore, the knockdown of USP28 efficiently decreased its target without influencing DTX3L mRNA expression (**Fig. S7B**). These data indicate that neither DTX3L nor USP28 regulates the other at the transcriptional level, suggesting that the mutual regulation of DTX3L and USP28 occurs at the level of protein stability.

As coupled changes in ubiquitination and protein amount are commonly the result of a changed half-life, we next measured whether DTX3L and USP28 could influence each otheŕs protein half-life after depletion of the one or the other with specific shRNAs in HEK293 cells. On the one hand, we found that the half-life of both DTX3L and USP28 is about 5 h when cells are transfected with scrambled shRNA **(Fig. 4A,C)**. As before, knockdown of DTX3L led to an increase in USP28 levels at time point zero of the cycloheximide challenge and further to an at least two-fold increase in USP28‘s half-life (**Fig. 4A,B**). On the other hand, we again observed that depletion of USP28 resulted in an increase of DTX3L levels at time zero and further also in an increase in it‘s half-life **(Fig. 4C,D)**. Similarly, depletion of USP28 in breast cancer cells (MDA-MB-231) also increased DTX3Ĺs half-life whereas overexpression of USP28 shortened the DTX3L half-life (**Fig. S8A,B**).

**Figure 4.**
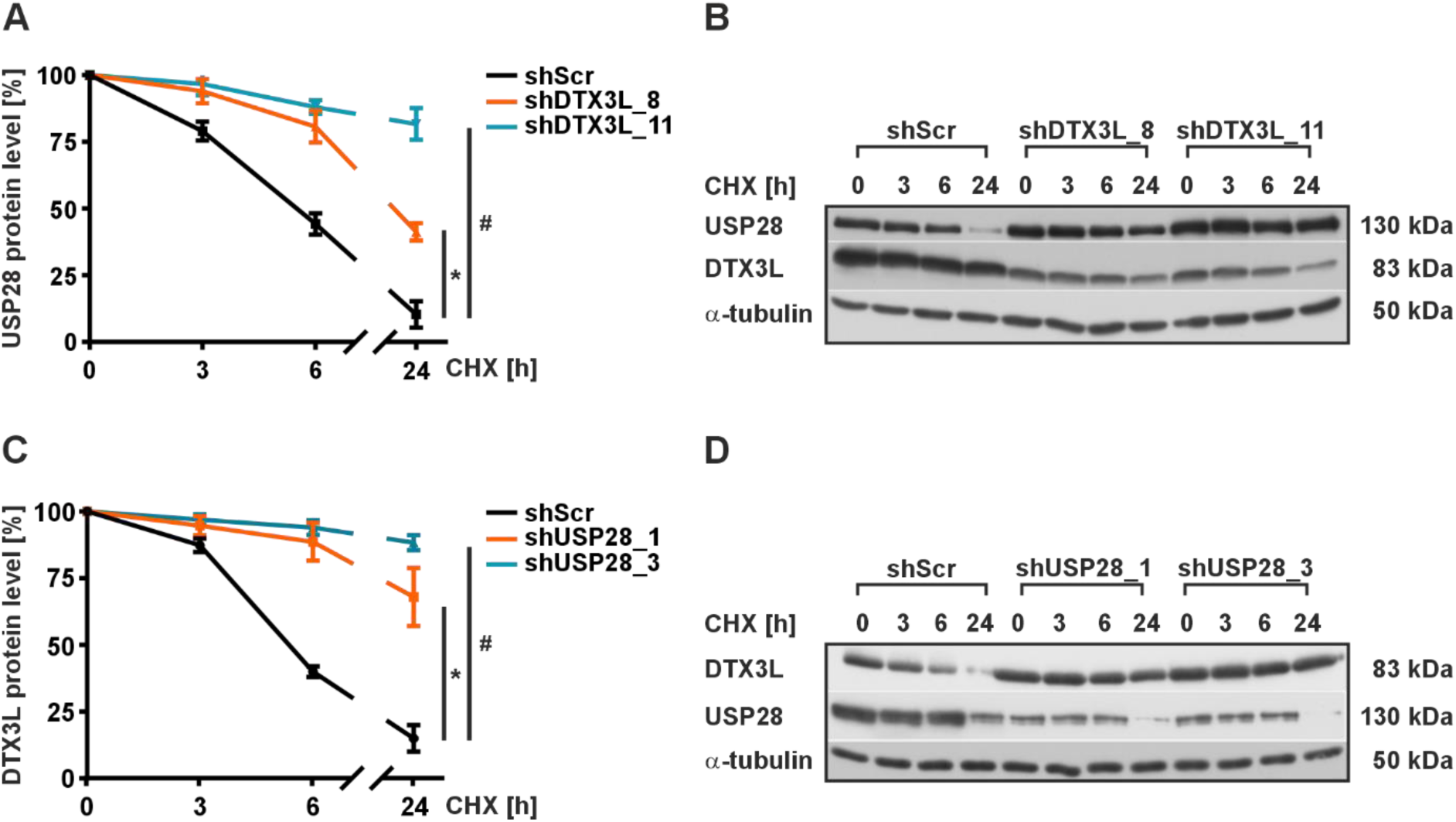
DTX3L and USP28 control each other’s half-life. **(A,C)** HEK293 cells were transfected with scrambled shRNA or with two different shRNÁs against either DTX3L or USP28. After transfection, protein synthesis was inhibited with cycloheximide (CHX; 10 µg/µl) and cells were harvested at indicated time points. In each experiment the protein levels at time point 0 were set to 100%. *significant differences shScr vs shDTX3L or shUSP28_1, ^#^significant differences shScr vs shDTX3L_11 or shUSP28_3, p ≤ 0.05. **(B,D)** Representative Western blot analysis. 100 µg of total protein lysate was analyzed with antibodies against USP28, DTX3L and α-tubulin.

To explore the degradation mechanisms for USP28 and DTX3L, we evaluated both proteasomal and lysosomal pathways by using MG132 as a proteasome inhibitor and chloroquine as a lysosome inhibitor. We found that the USP28-dependent reduction of DTX3L was effectively blocked in the presence of MG132 but not chloroquine (**Fig. S8C**). This suggests that USP28 facilitates the proteasomal degradation of DTX3L. Similarly, the effects of DTX3L were abolished by MG132 but not by chloroquine, indicating that the degradation of DTX3L is primarily proteasome-dependent (**Fig. S8D**).

Altogether, the current findings show that DTX3L and USP28 can mutually reduce each otheŕs half-life by involving the proteasome.

### DTX3L and USP28 control DNA repair pathways and affect cell viability

The most critical DNA lesions are double-strand-breaks (DSB). Several pathways are available to repair these detrimental lesions in cells. These include non-homologous end joining (NHEJ), homologous recombination (HR), single-strand annealing (SSA), and microhomology-mediated end joining (MMEJ). Canonical NHEJ directly joins both ends of the DSBs and is only partially error free ^27^, as a few nucleotide insertions or deletions may occur due to minor end processing. For comparison, HR uses homologous sequences of the sister chromatid as template and is therefore considered error-free. SSA is another example of homologous repair but requires extensive processing of the broken DNA ends before annealing between single stranded DNA repeats, resulting in deletions and therefore loss of genome integrity ^28^. MMEJ is different from SSA as it uses only microhomologies of a few base pairs between the two strands that arise after resection of the broken ends. As a consequence, MMEJ and SSA are always error-prone ^29^.

To analyze the participation of DTX3L and USP28 in DSB repair we induced genome-wide DSBs with bleomycin ^30^ and used an established EGFP-based reporter system allowing measurements of NHEJ, MMEJ, and both conservative HR and non-conservative SSA ^31–33^.

First, we examined whether the interaction between DTX3L and USP28 would be impaired after DSB induction, but the co-immunoprecipitations showed that this was not the case **(Fig. 5A)**. We then used the substrate EJ5SceGFP, which monitors all NHEJ events, that is, both simple rejoining between two cleaved I-*Sce*I sites as well as error-prone events accompanied by deletion or insertion of nucleotides ^33^. The results showed that USP28 knockdown reduced NHEJ frequencies by about 75%, whereas DTX3L knockdown increased it. Interestingly, the reduced NHEJ capacity in USP28 knockdown cells was rescued by simultaneous knockdown of DTX3L in these cells (**Fig. 5B,C,D**).

**Figure 5.**
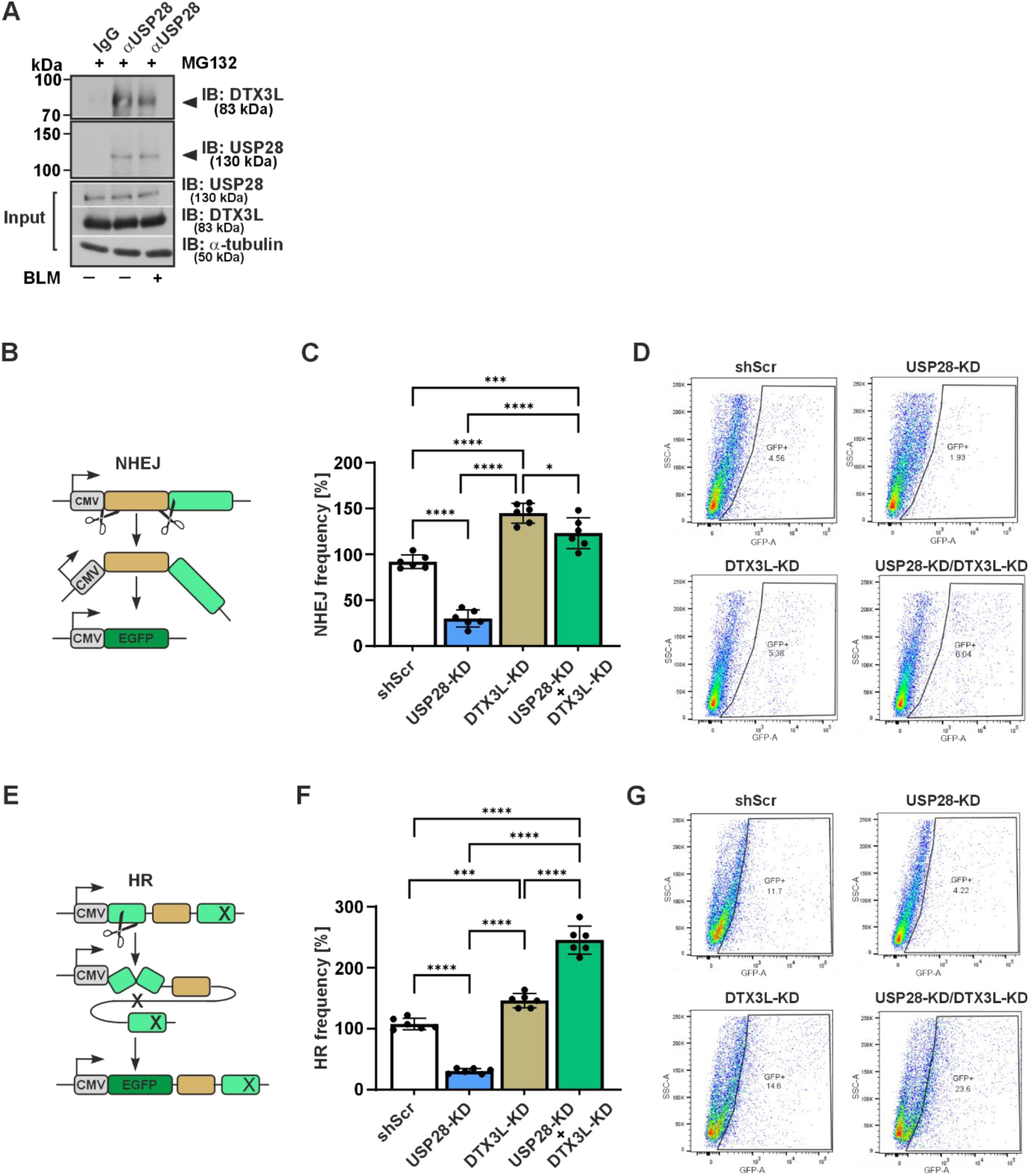
NHEJ-, and HR-DSB repair analysis. **(A)** Induction of DSBs by bleomycin (BLM) does not affect USP28/DTX3L interaction. Cells treated with BLM (10 µM) and the proteasome inhibitor MG132 were immunoprecipitated with either IgG control or USP28 antibodies. Resulting precipitates were analyzed by immunoblotting (IB) using a DTX3L antibody. Membrane was re-probed with an antibody against USP28. **(B, E)** Scheme of pathway-specific repair activities for NHEJ and HR measured with EGFP reporter assays. **(C, F)**. Quantification of NHEJ and HR DSB repair activities in BLM treated MDA-MB-231 Scr, USP28-KD, DTX3L-KD and USP28/DTX3L double knockdown cells. Mean values for the MDA-MB-231 Scr cells (shScr) were defined as 100%. Data are mean +/- SD from 6 measurements. The statistical significance of differences was determined using ordinary one-way ANOVA. **p<0.01, ****p<0.0001. **(D, G)** Representative flow cytometry images.

Next, we focused on HR using the substrate HR-EGFP/5′EGFP and found that knockdown of USP28 rendered the cells susceptible to DNA damage, indicated by the about 75% decrease in HR frequencies. After DTX3L knockdown HR frequencies were slightly increased. Importantly, the combined knockdown of USP28 and DTX3L induced HR frequencies by about 2-fold and the reduced HR capacity in USP28 knockdown cells was again rescued by knockdown of DTX3L **(Fig. 5E,F,G)**.

When SSA frequencies were evaluated, we again found that USP28 knockdown reduced SSA frequency by about 50%. Although DTX3L knockdown reduced SSA frequency by 20%, it counteracted the reduced SSA frequency seen in USP28 knockdown cells (**Fig. 6A,B,C**). Assessment of error-prone MMEJ using of EJ-EGFP ^31^ again showed that knockdown of USP28 strongly reduced MMEJ frequencies, whereas knockdown of DTX3L alone did not affect MMEJ frequencies. Again, the reduced MMEJ capacity in USP28 knockdown cells was rescued by simultaneous knockdown of DTX3L (**Fig. 6D,E,F**).

**Figure 6.**
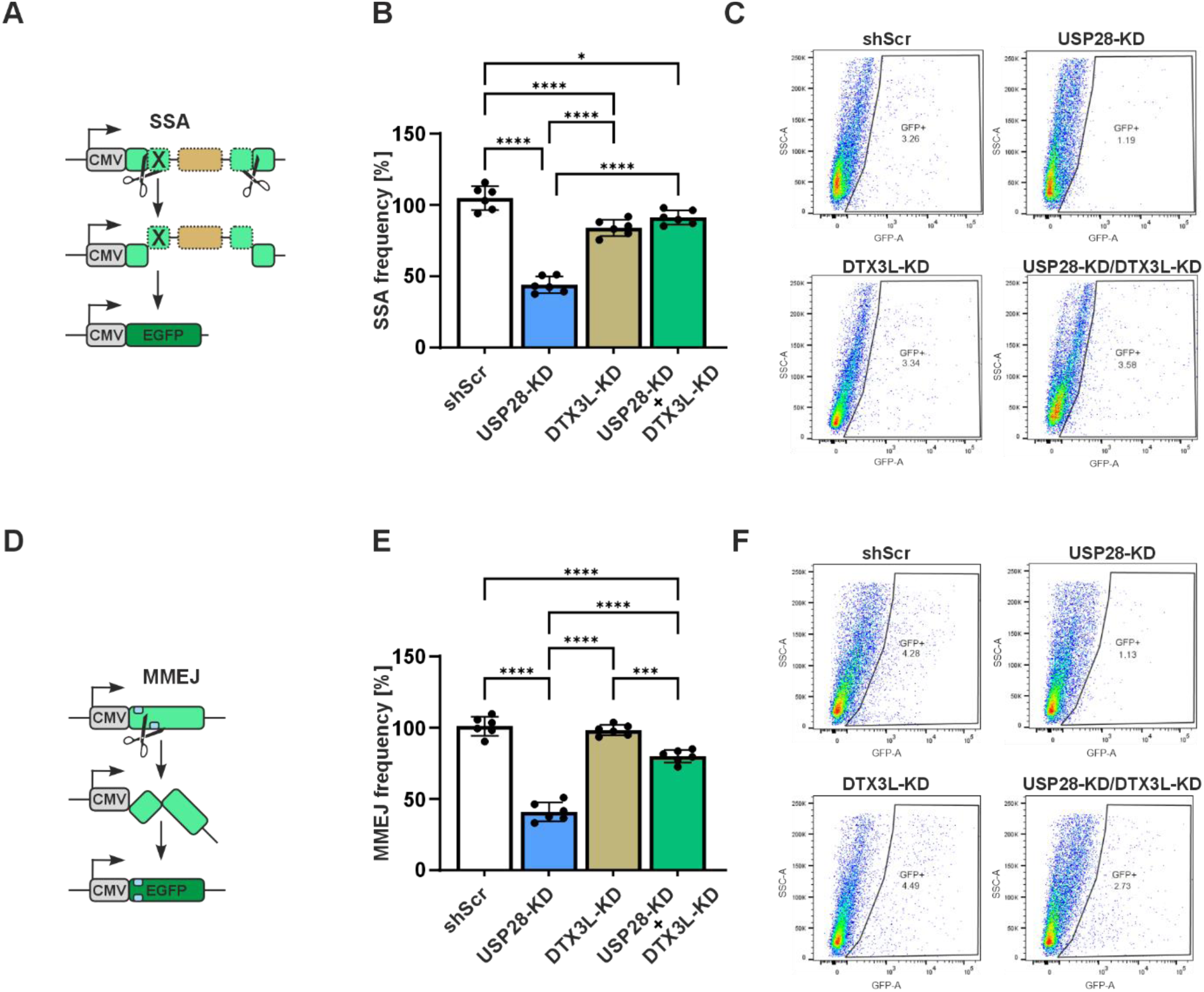
SSA-, and MMEJ-DSB repair analysis. **(A, D)** Scheme of pathway-specific repair activities for NHEJ and HR measured with EGFP reporter assays. **(B, E)**. Quantification of NHEJ and HR DSB repair activities in BLM treated MDA-MB-231 Scr, USP28-KD, DTX3L- KD and USP28/DTX3L double knockdown cells. Mean values for the MDA-MB-231 Scr cells (shScr) were defined as 100%. Data are mean +/- SD from 6 measurements. The statistical significance of differences was determined using ordinary one-way ANOVA. **p<0.01, ****p<0.0001. **(C, F)** Representative flow cytometry images.

Overall, these data show that silencing of USP28 reduces any DSB repair activity, which can be partially compensated by simultaneous silencing of DTX3L. Thus, it can be concluded that the regulatory interplay between USP28 and DTX3L modulates DSB repair. Since DNA repair mechanisms are crucial for maintaining cell viability, we investigated whether the loss of USP28 and DTX3L, alone or in combination, affects cell viability with potential pro-survival or pro-apoptotic outcomes in response to DSB induction. First, we analyzed the distribution of control and knockdown cells across different phases of the cell cycle. We found that single knockdown of USP28 had no or rather subtle effects on cell cycle distribution when compared to the control whereas knockdown of DTX3L resulted in a pronounced shift of cell populations into the G1/0 phase, thereby reducing the S phase and the G2 phase. Interestingly, the double knockdown cells showed a similar pattern except that they displayed a restored S phase (**Fig. 7A**). When assessing apoptosis, we found that USP28 knockdown increased the number of apoptotic cells, whereas DTX3L knockdown and double knockdown cells showed a similar pattern as the control group, indicating again that DTX3L can partially counteract the apoptotic effects caused by the USP28 knockdown alone (**Fig. 7B,C**). Taken together, our data suggest that USP28 and DTX3L may play antagonistic roles in regulating cell cycle progression and apoptosis by modulating the DNA damage response.

**Figure 7.**
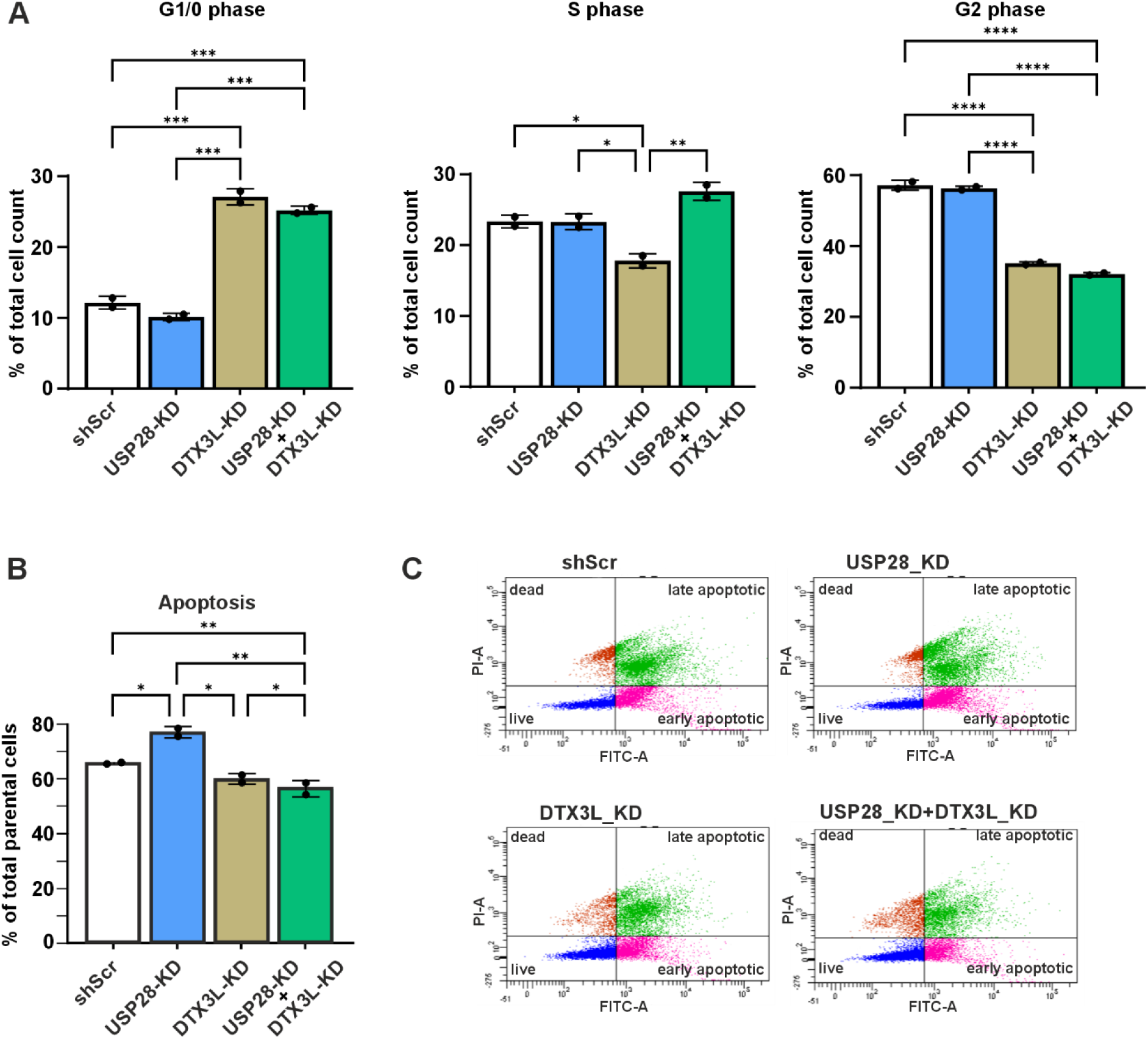
Mutual regulation of cell cycle and apoptosis by USP28 and DTX3L upon DNA DSB induction. MDA-MB-231 Scr, USP28-KD, DTX3L-KD and USP28/DTX3L double knockdown cells were treated with bleomycin (BLM; 10 µM) and further cultured for 24 h. **(A)** Cell cycle distribution of MDA-MB-231 shScr, USP28 KD, DTX3L KD and double knockdown (USP28 KD + DTX3L KD) cells in the G1/0 phase, S phase, and G2 phase. Significance * p<0.05, ** p<0.01, *** p<0.001 **(B)** Quantification of apoptosis in MDA-MB- 231 Scr, USP28 KD, DTX3L KD and USP28/DTX3L double knockdown cells Significance * p<0.05, ** p<0.01. **(C)** Histograms of apoptotic cells assessed by Annexin-V/PI staining and measured by flow cytometry.

## Discussion

The current study is the first one providing evidence for an interplay between the E3 ubiquitin ligase DTX3L and the DUB USP28. The findings present novelty in several aspects. First, the interplay between the two proteins is based on a direct physical transient interaction; second, they regulate each other’s levels in a mutual manner; and third, the reciprocal regulation affects USP28 target proteins and the cellular DSB repair capacity. These studies on the DTX3L– USP28 interaction suggest synergistic roles in DNA damage repair and cancer biology.

DTX3L was originally identified as a binding partner of PARP9, which is an oncogenic factor in diffuse large B-cell lymphoma (DLBCL) ^34^. Interestingly, those former studies indicated that DTX3L shuttles between the cytoplasm and the nucleus ^16^. Our current findings from co-immunoprecipitation experiments across cellular subfractions, supported by proximity ligation assays, confirm that both DTX3L and USP28 are present in both the cytoplasm and nucleus, where they interact with each other and may target specific proteins ^10,35^. Although cytoplasmic targets are still unknown, we found in our work that levels of HIF-1α, p53, and cMYC, all nuclear based transcription factors, were decreased upon forced expression of DTX3L and knockdown of USP28.

Preceding studies indicated that DTX3L is abundant at DNA lesion sites suggesting that it might be involved in the ADP-ribosylation dependent DNA damage response ^36,37^. More specifically, DTX3L in complex with PARP9 was shown to be recruited to sites of DNA damage by PARP1 ^9^. Different from PARP1, PARP9 was first described as a catalytically inactive member of the ARTD family ^38,39^ and more recently found to form a complex with DTX3L and regulate the E3 Ub ligase function of DTX3L ^5,8,40^. Simultaneously, PARP9 recognizes ADP-ribosylated proteins and therefore recruits the complexed E3 ligase to its substrate for ubiquitination and ultimately degradation ^41^. Given that USP28 is recruited to DNA damage sites by TP53BP1 ^15^ and that DTX3L affects the recruitment of TP53BP1 ^4^, we investigated the potential direct interaction between the Ub E3 ligase DTX3L and the DUB USP28, as well as the functional impact of their interaction ^4^.We purified full-length DTX3L and USP28 as well as its catalytically inactive mutant and studied ubiquitination in vitro and via immunoprecipitation in cellulo. These analyses demonstrated that DTX3L can ubiquitinate itself and that USP28 can remove the ubiquitin chains from DTX3L, which relied on the comparison of catalytically active (full-length USP28, catalytic domain of USP28) and inactive (USP28^C171A^) versions of USP28. Ablation of the DUB activity in USP28^C171A^ and the E3 ligase activity in DTX3L enabled us to demonstrate that also USP28 represents an ubiquitination substrate of DTX3L. Our MS measurements identified a wide distribution of Ub modified lysines along the USP28 molecule. Interestingly, K64, 85, 99, 115, and 135 were previously identified as sumoylation sites using an *in vitro* assay ^42^, and among these lysines, K99 was shown to be most efficiently sumoylated compared to others. These sumoylated and ubiquitinated amino acids reside within the N-terminus comprising the UBA domain and the SIM/UIMs. SUMO modifications in these regions negatively regulate USP28 catalytic activity^42^ and our results indicate that there could be a dynamic regulation of USP28 through competition between sumoylation and ubiquitination. All in all, these experiments revealed physical and functional interactions between DTX3L and USP28 assembling Ub E3 ligase and DUB activities, respectively, directed towards both enzyme components of the heterodimeric complex and to downstream proteins involved in signaling different types of stress such as hypoxia or DNA damage.

Although the involvement of DTX3L and USP28 in the regulation of the DNA damage response has been suggested by previous findings ^40^ ^19,20,43,44^, the current study provides much more detail on the direct interplay between DTX3L and USP28, particularly in the context of DSBs. USP28 appears to be essential for the efficient functioning of NHEJ, HR, SSA and MMEJ, as well as for cell survival after DNA damage, as shown by the severe impairment of DSB repair and subsequent increased apoptosis upon USP28 knockdown. Importantly, the partial rescue of the DNA repair pathways and the survival by simultaneous DTX3L knockdown suggests a compensatory mechanism and underscores the importance of their mutual interplay in maintaining genomic integrity. Apart from USP28 and DTX3L, this compensatory response may involve TP53BP1 and PARP9, both of which are known to be integral to DNA repair and cell cycle regulation ^20,31,45,46^. This is underscored by the fact, that we could co-precipitate TP53BP1, which is known to be stabilized by USP28 ^16^ together with DTX3L and USP28. Aside from TP53BP1, the compensatory mechanisms on the DNA damage response exerted by DTX3L and USP28 may involve NFκB signaling ^43^, which transcriptionally upregulates the NHEJ protein Ku70, the HR factors ATM and BRCA2 and activates BRCA1/CtIP ^47–50^.

The differential effects of DTX3L and USP28 knockdowns on cell cycle progression and apoptosis in response to DNA damage further elucidate their roles in the DNA stress response. Although USP28 knockdown did not alter the cell cycle compared to control cells, it promoted apoptosis, which is in line with the reduced DNA repair capacity. On the other hand, DTX3L knockdown appears to confer resistance to apoptosis, likely by mainly affecting the S phase of the cell cycle. This goes in line with the repair and senescence mechanisms activated by increased p53 which can be seen upon DTX3L knockdown, and which is in line with a previous study ^51^. Depending on the context or level of DNA damage, p53 can promote cell cycle arrest and DNA repair or apoptosis. In general, and as observed here, the arrest resulted in fewer cells entering the S phase which would allow the cells to repair the DNA damage ^52^. At the same time, p53 can induce cellular senescence ^53^, a state where cells remain metabolically active but do not proliferate. Those cells are less prone to apoptosis and do not progress through the cell cycle, contributing to the observed reduction in S phase and increased live cell counts.

Overall, our study provides essential insights into the molecular details of the cooperation between DTX3L and USP28. While the physical interactions highlight their role in catalyzing ubiquitination and deubiquitination processes, the functional interplay between DTX3L and USP28 reveals how they act like a rheostat, fine-tuning the cellular response to DNA damage and ultimately determining the fate of cells between survival and apoptosis. This balance underscores their multiple roles in the cell and sheds light on potential therapeutic targets to improve the DNA damage response in cancer treatment.

### Resource availability Lead contact

Requests for further information and resources should be directed to and will be fulfilled by the lead contact, Thomas Kietzmann.

### Materials availability

All unique/stable reagents generated in this study are available from the lead contact or co-authors with a completed materials transfer agreement.

### Data and code availability

Data are contained within the manuscript and the provided supporting information.

## Acknowledgements

We thank Philomena Schmid, Célia Tebbakh and Sabrina Arnold for technical assistance. The authors are grateful to Virpi Glumoff for FACS sorting with DNA repair GFP constructs. The use of the facilities and expertise of the Biocenter Oulu Structural Biology core facility (a member of Biocenter Finland, Instruct-ERIC Centre Finland and FINStruct), Proteomics and Protein Analysis core facility (a member of Biocenter Finland) and Biocenter Oulu sequencing center are gratefully acknowledged. MST experiments were carried out in the FINStruct site at the University of Helsinki. AZ1 was a generous gift from David M. Andrews (AstraZeneca).

## Author contributions

Y.A., L.L, T.K. conceptualization; Y.A, D.M., C.V.-R., H.I.A. and R.P.-H. investigation; M.R.-S and L.W. methodology; Y.A, D.M., C.V.-R., L.L., L.W. and T.K. writing-original draft; Y.A, D.M. C.V.-R., R.P.-R., M.R-S., L.W., L.L. and T.K. writing-reviewing and editing; L.L. and T.K. funding acquisition; R.P.-H., L.L. and T.K. supervision.

## Supplemental information

Figure S1-S11 and Tables S1.

## Methods

### Production of recombinant proteins

A codon optimised DNA encoding human USP28 was procured from Genscript and cloned between the NdeI/BamHI multiple cloning site (MCS) of pMJS162 ^54^. The expression vector for USP28 was then transformed into MDS42 bacterial cells and a starter culture was prepared in 5 ml of LB medium (Formedium, UK) with carbenicillin (100 µg/ml). After 14 h of incubation at 37°C, the starter culture was used to inoculate 500 mL of Terrific Broth (TB) autoinduction media with trace elements (Formedium,UK) supplemented with glycerol [0.8 (w/v)] and carbenicillin (100 µg/ml). Cultures were incubated briefly at 37°C until they reached and OD_600_ of 1.0, after which the temperature was decreased to 15°C for overnight incubation. Cells were harvested by centrifugation at 4200xg for 45 min at 4°C and re-suspended in lysis buffer [50 mM HEPES (pH 7.5), 500 mM NaCl, 0.5 mM TCEP, 10% (v/v) glycerol, 10 mM imidazole]. Re-suspended pellets were flash frozen in liquid N_2_ and stored at -20°C until purification.

To purify USP28, cell pellets were sonicated with a Branson digital sonifier using a ½ inch tip. The sonication was set for a total of 2.5 min of active sonication at a 40 % amplitude and cycles of 5 s of sonication with idling periods of 15 s. Lysates were centrifuged at 27,600×g at 4°C to recover soluble protein. The soluble fraction was filtered through a 0.45 µm sterile syringe filter and loaded onto a pre-equilibrated 5 mL HiTrap IMAC HP column (GE Healthcare Biosciences). The column was washed with 4 column volumes of lysis buffer followed by 4 column volumes of wash buffer 1 [50 mM HEPES (pH 7.5), 500 mM NaCl, 10% (v/v) glycerol, 0.5 mM TCEP, 50 mM imidazole]. To remove chaperone proteins, the column was washed with 4 column volumes of wash buffer 2 [50 mM HEPES (pH 7.5), 150 mM NaCl, 10% (v/v) glycerol, 10 mM imidazole, 0.5 mM TCEP, 5 mM ATP, 1 mM MgCl_2_]. Protein was recovered through a gradient elution of 60 mL from 10 mM imidazole to 500 mM imidazole. Pooled fractions from the elution were run through Superdex S200 16/600 gel filtration column with 150 mL of 30 mM HEPES (pH 7.5), 350 mM NaCl, 0.5 mM TCEP, 10% (v/v) glycerol, which also served as a storing buffer. Protein fractions were verified by SDS-PAGE and pooled, concentrated, flash frozen with liquid N_2_ and stored at –70°C.

DTX3L and other proteins used for ubiquitination reactions were purified as described previously ^8^. Nucleic acid contamination was assessed using absorbance ratio of 260 nm over 280 nm with ratios of 0.7 for DTX3L and 0.5 for USP28.

### *In vitro* ubiquitination assays

For the *in vitro* deubiquitination, we prepared 50 µl reactions. Proteins were diluted in 50 mM Hepes pH 7.5, 50 mM NaCl. Protein concentrations used were Ub (50 µM), E1 (0.4 µM), UbcH5a (E2) (2 µM) and DTX3L (0.7 µM). USP28 was used at a final concentration of 3 µM. Reactions were assembled on ice and then initiated by addition of ubiquitination buffer (20X concentration: 1M tris pH 7.5, 40 mM ATP, 100 mM MgCl2, 40 mM DTT) to a final concentration of 1X and incubated at room temperature for 2-3 h. Outcomes of the reactions were analyzed by adding 15 µL of the reaction to an SDS-PA gel, which was run for 45 min at 200 V and then proteins were visualized with either Coomassie blue staining or by Western blotting. Blotting of ubiquitination reactions was done by wet tank transfer to a nitrocellulose membrane at 100 V for 1 h. Time dependent ubiquitination reactions were performed in the same manner, except that the total reaction volume was 40 µL. Ubiquitination sites in USP28^C171A^ were identified as described in ^8^.

### Microscale thermophoresis (MST)

The dissociation constant (K_d_) between DTX3L and USP28 was measured with Monolith (NanoTemper) using a RED-NHS 2^nd^generation labeling dye (NanoTemper). NHS ester groups covalently crosslink lysine residues. DTX3L was labeled by adding 30 µM dye to 10 µM protein, 100 µl of ligand buffer [20 mM HEPES (pH 7.4), 350 mM NaCl, 0.5 mM TCEP]. The labeling reaction was incubated for 60 min at room temperature, in the dark, and excess dye was removed with a desalting column pre-equilibrated with 12 ml of ligand buffer. Labeled protein was recovered in 450 µL of ligand buffer. The degree of labeling was determined to be 90% based on the absorbances at 280 nm and 650 nm.

For affinity measurements, labelled DTX3L (20 nM final concentration) was added to a serial dilution of USP28 (22.5 µM to 0.68 nM). Immediately after mixing, 10 µl were loaded onto the Monolith NT.115 Premium Capillaries (NanoTemper). The MST signal was recorded with NT.Control v2.1.31 software and data were fitted to a 4-parameter sigmoidal curve using Graphpad Prism version 9.3.1. (GraphPad Software, La Jolla, CA, USA). Three independent MST measurements were performed.

### Cell cultures

SK-MES-1, human embryonic kidney 293 (HEK 293) and HeLa cells were cultured under normoxia (16% O_2_, 79% N_2_ and 5% CO_2_ [by volume]) in minimal essential medium (MEM) supplemented with 10% fetal bovine serum (FBS). PC-3 cells were maintained in RPMI1640 medium supplemented with 10% FBS. MDA-MB-231 cells with stable USP28 knockdown were generated as described ^23^ and cultured in DMEM supplemented with 10% FBS.

All cell lines were tested Mycoplasma negative by using the MycoAlert Detection Kit (Lonza). In all experiments the number of cell passages used was below 10. For protein extraction, cells were seeded onto 6 cm dishes. After a medium change, cells were treated with the USP25/28 inhibitor AZ1 (10 µM), further cultured for 24 hours either under normoxia or hypoxia (5% O_2_, 90% N_2_, and 5% CO_2_) and then harvested. In all experiments control cells were treated with DMSO.

### Plasmid constructs for knockdown and protein expression

The constructs for pDZ-Flag-USP28, pRetrosuper-USP28 shRNA-1, and pRetrosuper- USP28 shRNA-3 were described previously ^21^. The full length USP28 construct with mutation in cysteine 171 was generated using the QuickChange mutagenesis kit (Promega) as described ^21^.

The constructs for pLKO.1-shDTX3L_8 (TRCN0000073208) and pLKO.1-shDTX3L_11 (TRCN0000073211) were from Sigma Aldrich MISSION shRNA library, distributed by Genome Biology Unit core facility supported by HiLIFE and the Faculty of Medicine, University of Helsinki, and Biocenter Finland.

The constructs for pRK5-HA-Ubiquitin-WT (Addgene plasmid #17608), pRK5-HA-Ubiquitin-KO (Addgene plasmid #17603), pRK5-HA-Ubiquitin-K48 (Addgene plasmid #17605) and pRK5-HA-Ubiquitin-K63 (Addgene plasmid #17606) were a kind gift from Ted Dawson and described previously ^55^.

The construct for DTX3L^CS^ was obtained through site-directed mutagenesis as previously described ^56^. DTX3L deletion mutants were obtained through sequence and ligation independent cloning (SLIC) in pcDNA3.1 HA vector from PCR amplification of the D1-D3 domains (A2-A516) and RING-DTC domains (K555-E740).

### Western blot analysis, protein half-life studies and co-immunoprecipitation

Western blot analysis was carried out as described previously ^57^. In brief, purified proteins or total mammalian cell lysates were separated by sodium dodecyl sulfate (SDS)-polyacrylamide gel electrophoresis. After electrophoresis and electroblotting onto a nitrocellulose membrane, membranes were blocked at room temperature for 1 h with 1X casein blocking solution (Bio-Rad) (#1610782) or 5% milk powder in 1X PBS-Tween. Proteins were detected with antibodies shown in Supplementary Table S1. The ECL system (GE Healthcare, Germany) was used for detection. Blots of three independent experiments were quantified by densitometry with the Image Quant TL Program (GE Healthcare); densitometry data were normalized to α-tubulin.

For half-life studies, HEK 293 cells were transfected with expression vectors encoding full length USP28, USP28^C171A^ or full length DTX3L. After 24 h, cycloheximide (10 µg/ml; Sigma-Aldrich) was added to the medium, cells were harvested at the indicated time points and protein levels were measured by immunoblot analysis. Blots of three independent experiments were quantified by densitometry with the Image Quant TL Program (GE Healthcare); densitometry data were normalized to α-tubulin.

For co-immunoprecipitation, the cells were pretreated with the proteasome inhibitor MG 132 (50 µM; Calbiochem) for 4 h. Cells were washed twice with ice-cold 1x PBS, then (dithiobis(succinimidyl propionate)) DSP was added to a final concentration of 2 mM and cells incubated for 2 h at 4°C. To terminate the reaction, glycine was added to a final concentration of 10 mM for additional 15 minutes. Then cells were scraped in lysis buffer (50 mM Tris/HCl, pH 7.5, 150 mM NaCl, 1% Triton X-100, 2 mM EDTA, 2 mM EGTA, 1 mM PMSF, and complete protease inhibitor cocktail tablet; Roche), incubated with continuous shaking at 4°C for 20 min and then centrifuged at 12 000xg at 4°C for 15 min. Cytoplasmic and nuclear fractions from PC-3 cells were isolated using a nuclear extraction kit (#2900; Merck Millipore) according to the manufactureŕs protocol. To recover immunoprecipitates, 150 µg of protein lysate was incubated with 2 µg of antibodies against TP53BP1, USP28, DTX3L, HA and FlagM2 epitope for 1 h at 4°C before Protein G Sepharose beads (30 µL per reaction mixture; GE Healthcare #GE17-0618-01) were added for 12 h. Thereafter, the beads were washed 5 times with lysis buffer and recovered, pellets were dissolved in 2x Laemmli buffer, loaded onto a 7.5% SDS gel, blotted, and detected with Abs against DTX3L, USP28 and ubiquitin.

### Proximity ligation assay

DTX3L-USP28 interaction in SK-MES-1 cells was detected with Duolink PLA Kit (Olink Bioscience, Sweden; PLA probe anti-rabbit plus. PLA probe anti-mouse minus, Detection kit orange, # 92002-0030) according to the manufactureŕs protocol. Briefly, cells were grown on coverslips until they reached sufficient confluency and fixed with 4% paraformaldehyde. Then cells were permeabilized with blocking buffer and incubated with primary rabbit polyclonal USP28 antibody (1:100; #HPA006778; Sigma-Aldrich) and mouse monoclonal DTX3L antibody (1:100, #sc-514776, Santa Cruz) diluted in blocking solution over night at 4°C. In the negative control DTX3L antibody was omitted. After the last washing step with buffer B, cells were stained with bisBenzimidine (1:5000; Hoechst 33342; #14533; Sigma-Aldrich) for 5 min and mounted with Immu-Mount (#9990402; Fisher Scientific). Microscopy was performed using a confocal microscope (LSM700, Zeiss).

### RNA preparation, reverse transcription and quantitative real-time PCR

Isolation of total RNA was performed using RNeasy Mini Kit (Qiagen, Hilden, Germany) following the manufactureŕs protocol. Total RNA concentration and purity were measured on a NanoDrop spectrophotometer. 1 µg of total RNA was used for cDNA synthesis using iScript cDNA Synthesis Kit (Bio-Rad, München, Germany).

Quantitative real-time PCR was performed using iTaq SYBR Green Supermix (Bio-Rad) in a CFX96 Touch Real-Time PCR Detection System (Bio-Rad) with the primers shown in Supplementary Table S2. The experiments for each data point were carried out in triplicate. The relative quantification of gene expression was determined using the ΔΔCt method ^58^.

### Immunofluorescence

MDA-MB-231 cells were grown on coverslips. When cells reached sufficient confluency, media was removed, cells were washed with PBS and fixed in 4% paraformaldehyde for 10 min and washed with PBS. Blocking was performed in PBS containing 5% fetal bovine and 5% goat serum for 1 h. This was followed by incubation with primary antibodies against DTX3L (1:100, #sc-514776, Santa Cruz) and USP28 (1:100; #HPA006778; Sigma-Aldrich); both were diluted in 1% blocking buffer and incubated with the coverslips overnight at 4°C. After overnight incubation samples were washed in PBS and incubated with Alexa Fluor anti-mouse 488 (#A-11001; Invitrogen) and anti-rabbit 546 (#A-11035; Invitrogen) secondary antibodies diluted in 1% blocking buffer at room temperature for 1 h, respectively. Cell nuclei were visualized by Hoechst 33342 staining (#14533; Sigma-Aldrich). The specimens were mounted with Immu-Mount (#9990402; Fisher Scientific) and fluorescent images were obtained by confocal microscopy (LSM700, Zeiss).

### DSB repair

Analysis of DSB repair was conducted as described ^31,33,59^. In brief, plasmid constructs expressing the endonuclease I-*Sce*I (pCMV-I-SceI) and reporter constructs encoding different DSB repair substrates for NHEJ (EJ5SceGFP), MMEJ (EJ-EGFP), homologous recombination, HR (HR-EGFP/5’EGFP), and SSA (5’EGFP/HR-EGFP) were introduced by nucleofection according to the Amaxa protocol (Lonza, Cologne, Germany). To balance the DNA amount and to control transfection efficiencies, transfection mixtures for split cell culture samples contained either pBlueScriptII KS (pBS, Stratagene, Heidelberg, Germany) or wild-type EGFP expression plasmid, respectively. Mean transfection efficiencies were in the range of 50%. After transfection, cells were cultured for 24 h, then treated with bleomycin (10 µM) and further cultured for additional 24 hours. Thereafter, DSB repair was monitored via quantification of EGFP-positive cell fractions with a LSR Fortessa (Becton Dickinson) flow cytometer and analyzed by the diagonal gating method in the Fl1/Fl2 dot plot as described ^31^ by using the FlowJo 10.7.1. program. Each quantification of green fluorescent cells in repair assays was normalized by use of the individually determined transfection efficiency to calculate the DSB repair frequency Negative and positive controls for transfection efficiency are provided in **Fig. S11**.

### Analysis of cell cycle and cell death

Cell cycle and cell apoptosis were measured in synchronized cells by Annexin-V-FLOUS and propidium iodide (PI) staining kits (Roche/Sigma-Aldrich, Helsinki, Finland) according to the manufacturer‘s protocol. For synchronization, confluent cells were kept without medium change for 24 h. Release into the cell cycle was achieved by subculture and incubation with fresh serum with or without bleomycin (10 µM) for 24 hours. Apoptosis and cell cycle parameters were analyzed by flow cytometry with a LSR Fortessa (Becton Dickinson).

### Statistical Analysis

Densitometry data were plotted as fold induction of relative density units, with the zero-value absorbance in each figure set arbitrarily to 1 or 100%. If not otherwise stated statistical comparisons of absorbance differences were performed by the Mann-Whitney test (Statview 4.5, Abacus Concepts, Berkeley, CA), and p values p < 0.05 were considered significant.

Due to the non-saturating conditions of the experiment, the estimation of the K_d_ was done by fitting the data in logarithmic scale using a four-parameter logistic regression model (4PL) in GraphPad Prism version 9.3.1. Averages were taken from the log(Kd) values to allow reporting of a symmetric standard deviation.

