## Supplementary information for "The E3 ubiquitin ligase DTX3L and the deubiquitinase USP28 fine-tune DNA double strand repair through mutual regulation of their protein levels"

### CONTENTS

**Figure S1.** DTX3L and USP28 colocalize and interact with each other.

**Figure S2.** CD spectra of USP28 and the inactive mutant.

**Figure S3.** MST binding curve for DTX3L and USP28.

**Figure S4.** MST traces.

**Figure S5.** Hydrolysis of pre-formed poly-ubiquitin chains by USP28.

**Figure S6.** Ubiquitination reactions and CO-IP with a DTX3L inactive mutant ($DTX3L^{CS}$)

**Figure S7.** Mutual regulation of DTX3L and USP28 protein levels.

**Figure S8.** Mutual regulation of USP28 and DTX3L protein levels by proteasomal degradation.

**Figure S9.** FACS analysis of DNA DSB repair activity by USP28 and DTX3L

**Figure S10.** Regulation of DNA DSB repair activity by USP28 and DTX3L

**Figure S11.** Controls for flow cytometry.

**Table S1.** Antibodies used in the study.

**Table S2.** Primers used in the qRT-PCR analyses.

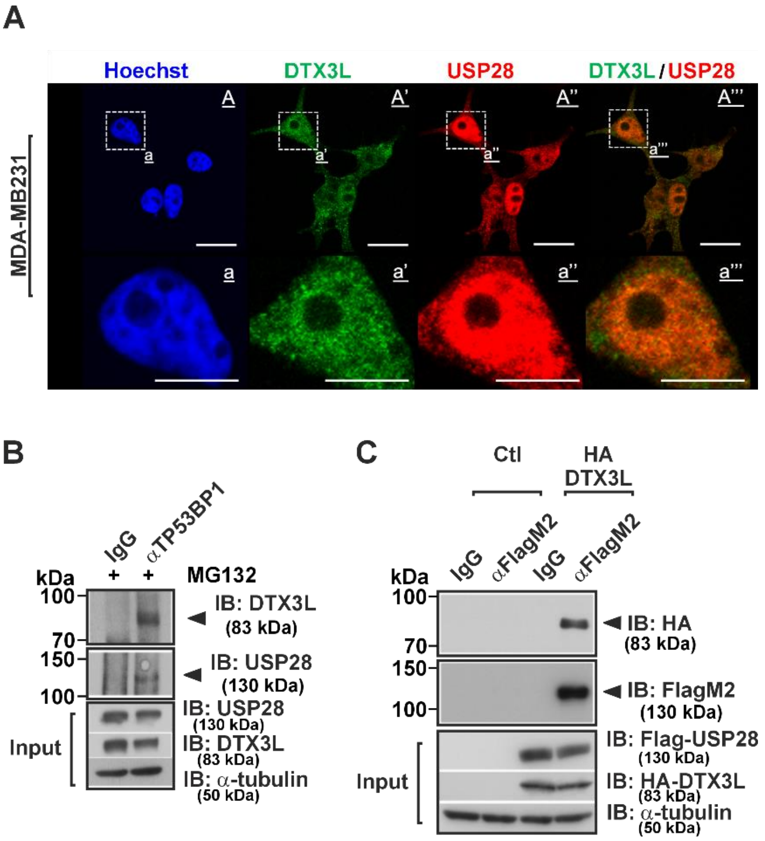


**Figure S1.** DTX3L and USP28 colocalize and interact with each other. **(A)** Immunostaining with DTX3L (green) and USP28 (red) antibodies in MDA-MB-231 cells shows that both proteins are co-expressed in the same cellular compartments. DAPI stained nuclei shown in blue. Scale bars: A-A''' 20 µm and a-a''' 10 µm. **(B)** SK-MES-1 cells were treated with the proteasome inhibitor MG132 (50 µM) for 4 hours. Anti-TP53BP1 immunoprecipitates (IP) were analyzed by immunoblotting (IB) with DTX3L or USP28 antibodies, respectively. **(C)** HEK293 cells were transfected with empty control vector (Ctl) or expression vectors encoding HA-tagged DTX3L and Flag-tagged USP28. Cell extracts were immunoprecipitated with either IgG control or FlagM2-tag antibodies. Resulting precipitates were first analyzed by immunoblotting (IB) using a HA-tag antibody to detect DTX3L; thereafter, the membrane was re-probed with a FlagM2 antibody to detect USP28.

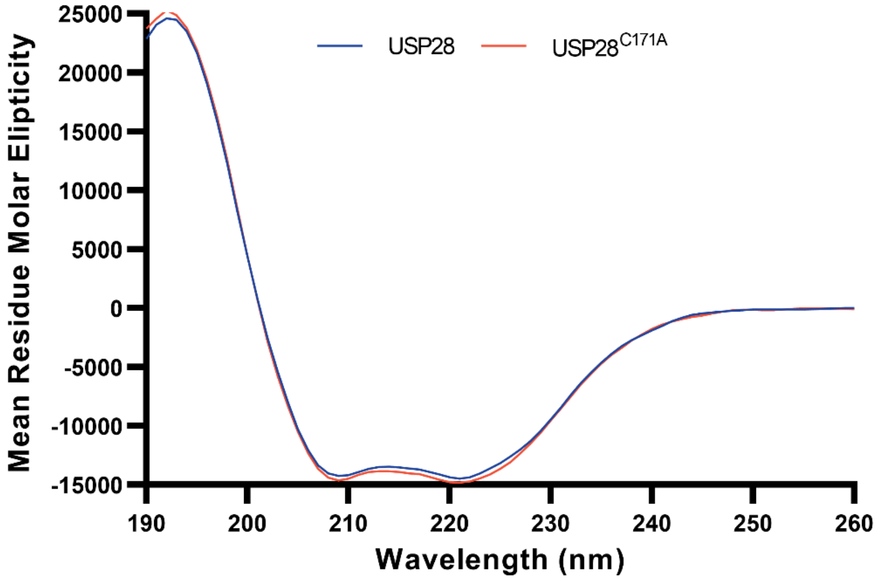


**Figure S2**. CD spectra for USP28 and USP28C171A. The spectra correspond to properly folded proteins indicating that the mutation does not affect the folding status.

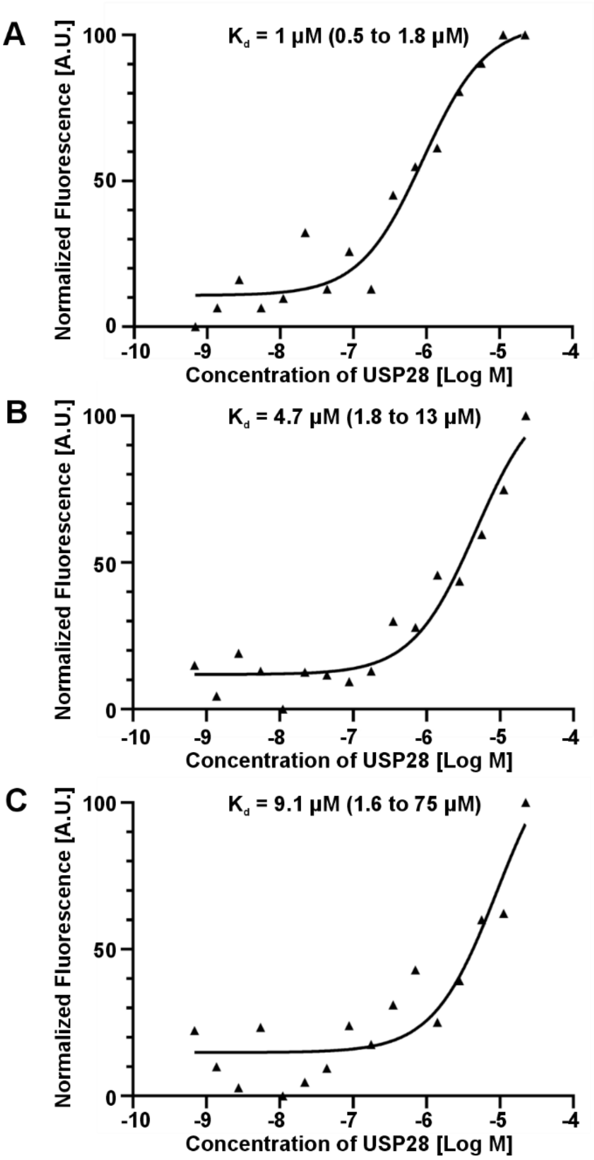


**Figure S3.** MST binding curves for DTX3L and USP28**.** The curves represent individual experiments at constant concentration of DTX3L and varying concentrations of USP28. DTX3L was labelled with RED-NHS 2$^{nd}$ generation labelling (NanoTemper) dye for 60 min at room temperature and excess dye was removed with a desalting column before the measurements. Data was fitted to a $K_d$ model with MO.Affinity Analysis (NanoTemper). The determined $K_d$ for each curve is marked on top with the confidence interval stated between brackets.

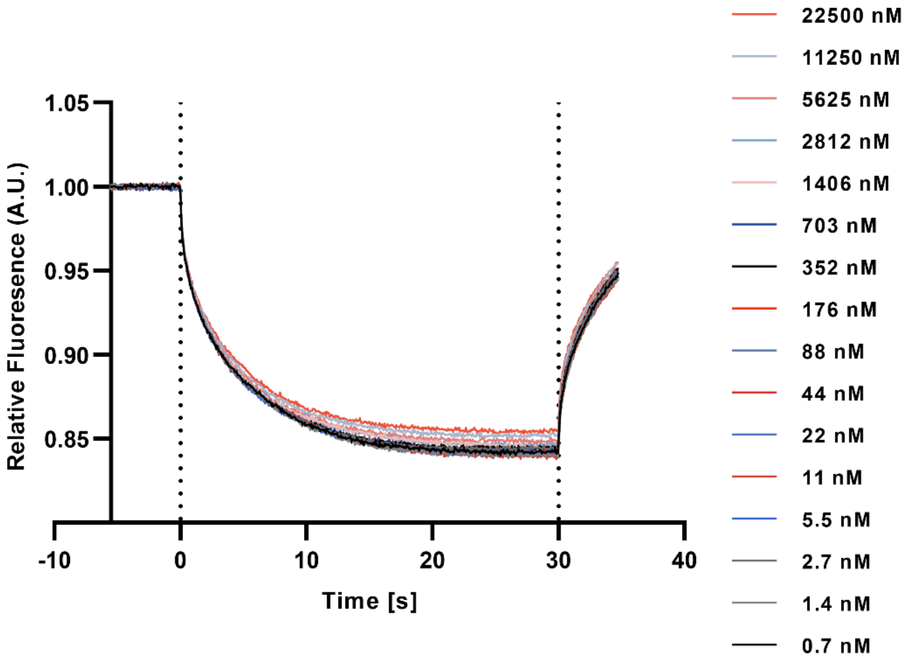


**Figure S4**. MST trace for DTX3L with different concentrations of USP28. Binding curve (Figure 1C) was determined from the relative fluorescence as a function of time. A stable reading before the start of the thermophoresis (up to 0 s) indicates that an increase in the fluorescence is not likely caused by unspecific binding.

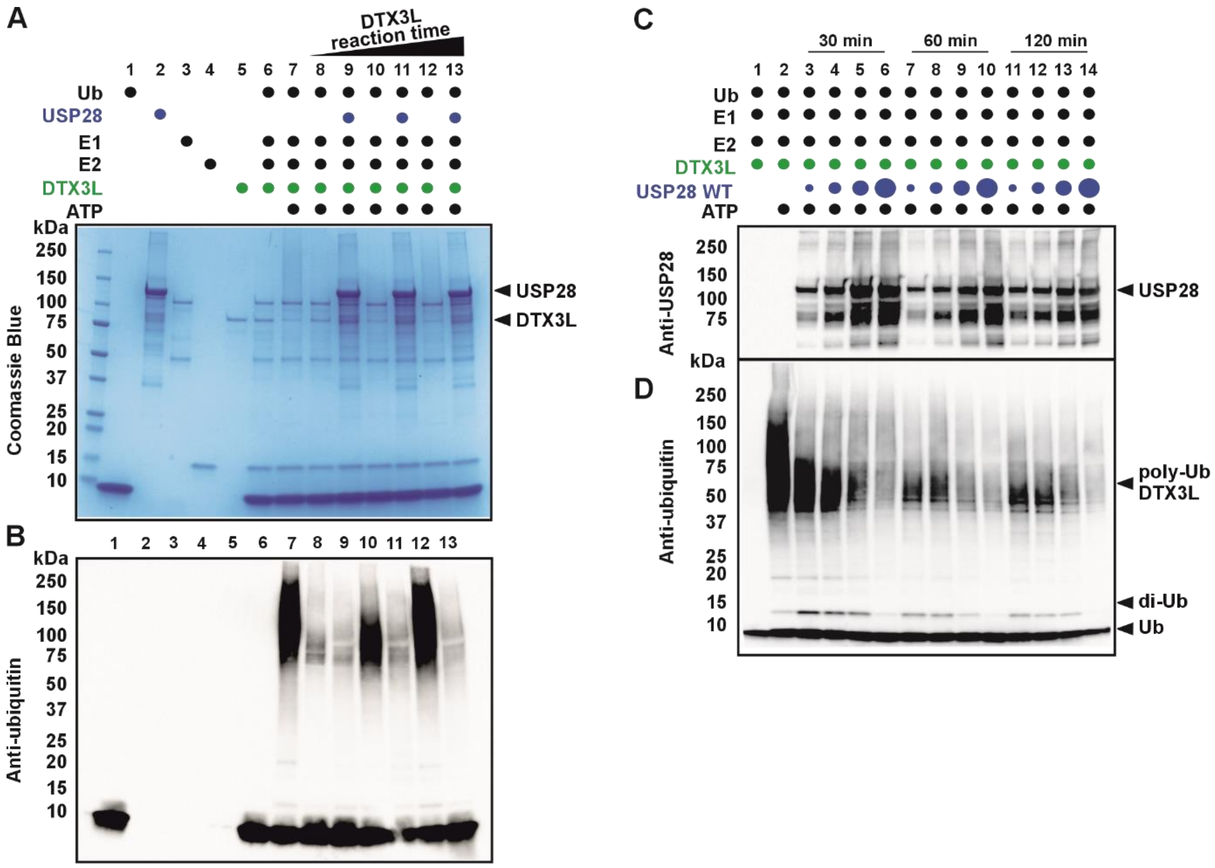


**Figure S5.** Hydrolysis of pre-formed poly-ubiquitin chains by USP28 is a time- and concentration-dependent event. Ubiquitination reactions were conducted by mixing ubiquitin (45 µM), Ube1 (0.4 µM), Ube2D1 (2 µM) and DTX3L (1 µM) in 40 µL of 50 mM HEPES (pH 7.5), 50 mM NaCl. The reaction was initiated by the addition of ubiquitination buffer [20x concentration: 1 M Tris (pH 7.5), 40 mM ATP, 100 mM MgCl2, 40 mM DTT] to a final concentration of 1x. To assess the hydrolysis of pre-formed poly-ubiquitin chains **(A and B)**, reactions were incubated for 30, 60 and 120 minutes and subsequently treated with USP28 (3 µM) (**A and B, lanes 9, 11 and 13**). The time- and concentration-dependency of the hydrolysis **(C and D)** was assessed by adding USP28 (0.5, 1, 2 or 3 µM) to ubiquitination reactions that have been incubated for 2 h. To stop the hydrolysis, 4x Laemmli buffer was added after 30, 60 and 120 min. The increasing concentration of USP28 in the **C and D** is represented by an increase in size of blue dots. Ubiquitination buffer was omitted in the negative control reaction (**A and B lane 6, and C and D lane 1**). The omission of USP28 served as a ubiquitination positive control (**A and B lane 7, and C and D lane 2**). The reactions were analysed by SDS-PAGE Coomassie blue staining **(A)** or by Western blotting with anti-ubiquitin antibody **(B and D)** or anti-USP28 antibody **(C)**.

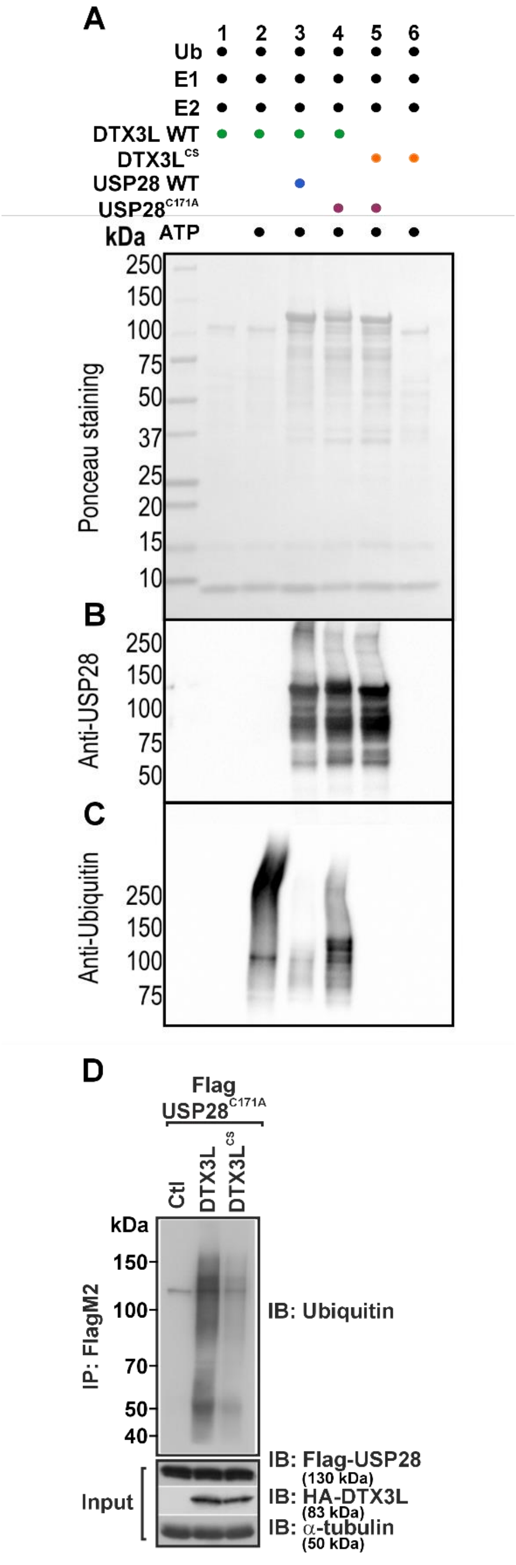


**Figure S6.** Ubiquitination reactions and CO-IP with a DTX3L E3 ligase inactive mutant (DTX3L$^{CS}$). Ubiquitination reactions were conducted by mixing ubiquitin (20 µM), Ube1 (0.4 µM), Ube2D1 (2 µM), DTX3L or DTX3L$^{CS}$ (1 µM), and USP28 or USP28$^{C171A}$ (3 µM) in 50 µL of 50 mM HEPES (pH 7.5), 50 mM NaCl. The reaction was initiated by the addition of ubiquitination buffer [20x concentration: 1 M Tris (pH 7.5), 40 mM ATP, 100 mM $MgCl_2$, 40 mM DTT] to a final concentration of 1x. Ubiquitination buffer was omitted in the negative control reaction (lane 1). Lane 2 served as positive control. The reactions were incubated for 4 hours and stopped with 4x Laemmli buffer. Lane 3 shows the hydrolysis activity of USP28. The modification of USP28$^{C171A}$ by DTX3L is shown in lane 4. The lack of ubiquitination activity of DTX3L$^{CS}$ on a substrate or on itself is shown in lanes 5 and 6, respectively. The reactions were analysed by Ponceau staining **(A)** or by Western blotting with anti-USP28 **(B)** and anti-Ub antibodies **(C)**. **(D)** HEK293 cells were cotransfected with the catalytic inactive FlagM2-USP28 (USP28$^{C171A}$) mutant and an empty control vector (Ctl) or expression vectors encoding HA-tagged DTX3L WT or with the catalytic inactive HA-tagged DTX3L (DTX3L$^{CS}$) mutant. Cell extracts were immunoprecipitated with either IgG control or FlagM2-tag antibodies. Resulting precipitates were analyzed by immunoblotting (IB) using an ubiquitin antibody to detect DTX3L ubiquitinylation.

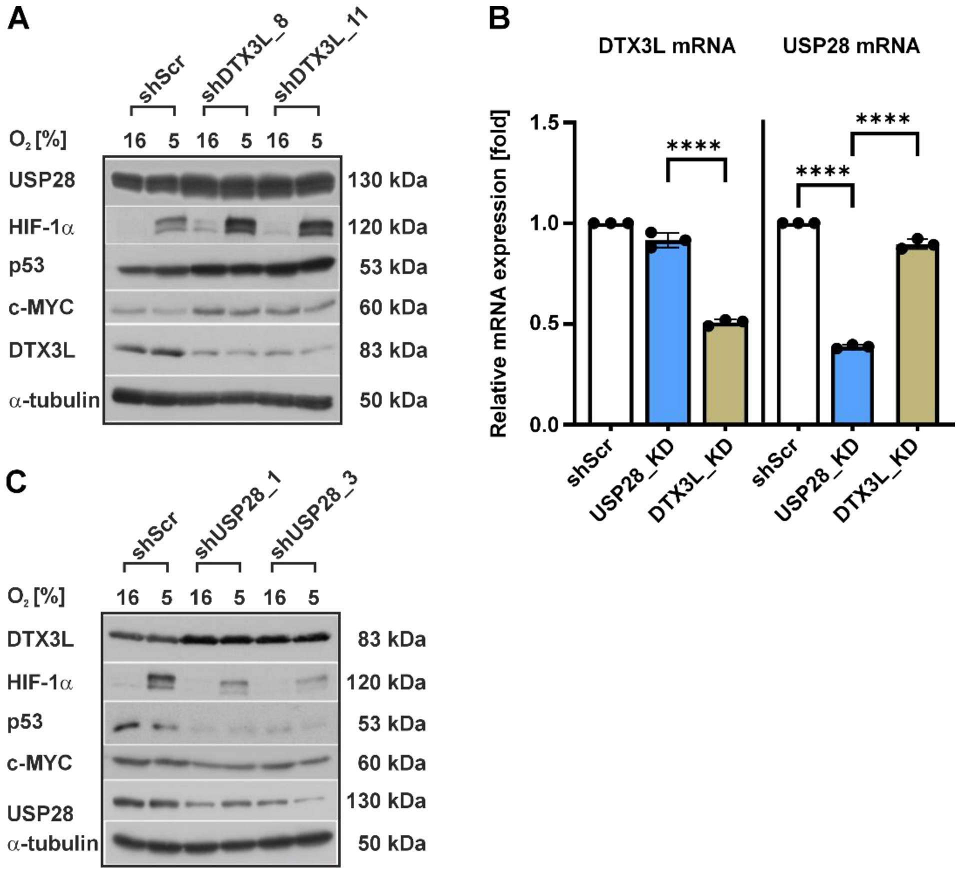


**Figure S7.** Mutual regulation of DTX3L and USP28 protein levels. **(A,C)** HeLa cells were transfected either with an expression vector for scrambled control shRNA (shScr), or one of two independent shRNAs (shDTX3L_8, shDTX3L_11) against DTX3L or one of two independent shRNAs (shUSP28_1, shUSP28_3) against USP28. After transfection, cells were further cultured under normoxia (16% $O_2$) or hypoxia (5% $O_2$) for 4 hours. USP28, HIF-1α, p53, c-MYC, and DTX3L protein levels were measured by Western blot analysis. Alpha tubulin served as a loading control. **(B)** SK-MES-1 cells were transfected either with an expression vector for scrambled control shRNA (shScr), or in USP28_KD with shRNA_1 against USP28 or in DTX3L_KD with shDTX3L_8. After transfection, cells were further cultured under normoxia or hypoxia for 6 h. DTX3L and USP28 mRNA levels were measured by qRT-PCR. Mean values for shScr cells were defined as 1. Data are mean +/- SD from 3 measurements. The statistical significance of differences was determined using ordinary one-way ANOVA. ****p<0.0001.

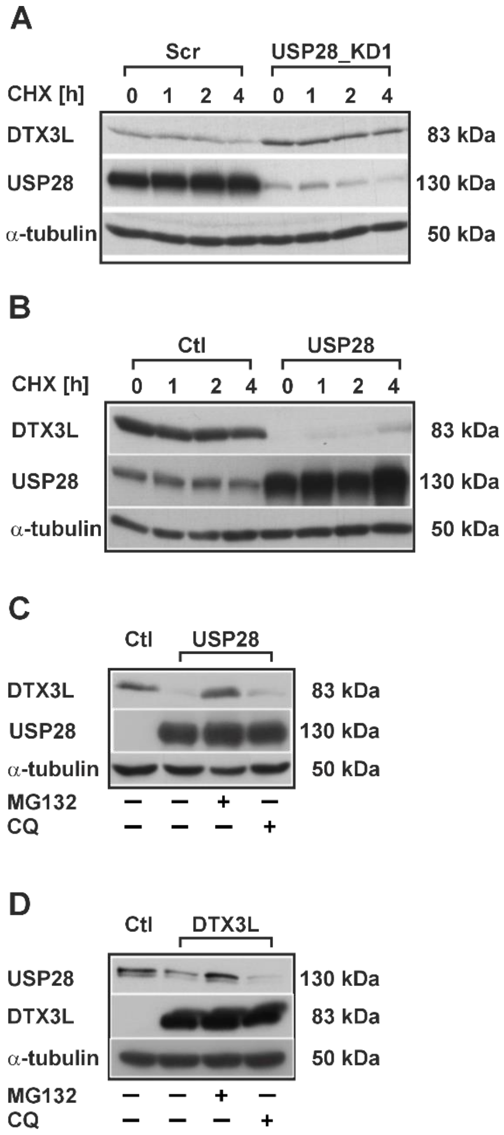


**Figure S8. Mutual regulation of USP28 and DTX3L protein levels by proteasomal degradation (A)** Protein synthesis in MDA-MB231 Scr and USP28 KD1 cells was inhibited by cycloheximide treatment (CHX, 10 µg/ml) and DTX3L protein half-life was assessed by Western blot analysis. Scr, scrambled; KD, knockdown. HEK293 cells were transfected with empty vector control or expression plasmids encoding either for full-length USP28 or full length DTX3L. **(B)** After transfection, protein synthesis was inhibited with cycloheximide (10 µg/ml) and DTX3L protein half-life assessed by Western blot analysis. **(C,D)** After transfection, cells were treated with the proteasomal inhibitor MG132 (50 µM) for 4 hours or with the lysosomal inhibitor chloroquine (CQ; 50 µM) for 24 h. USP28 and DTX3L protein levels were measured by Western blot analysis.

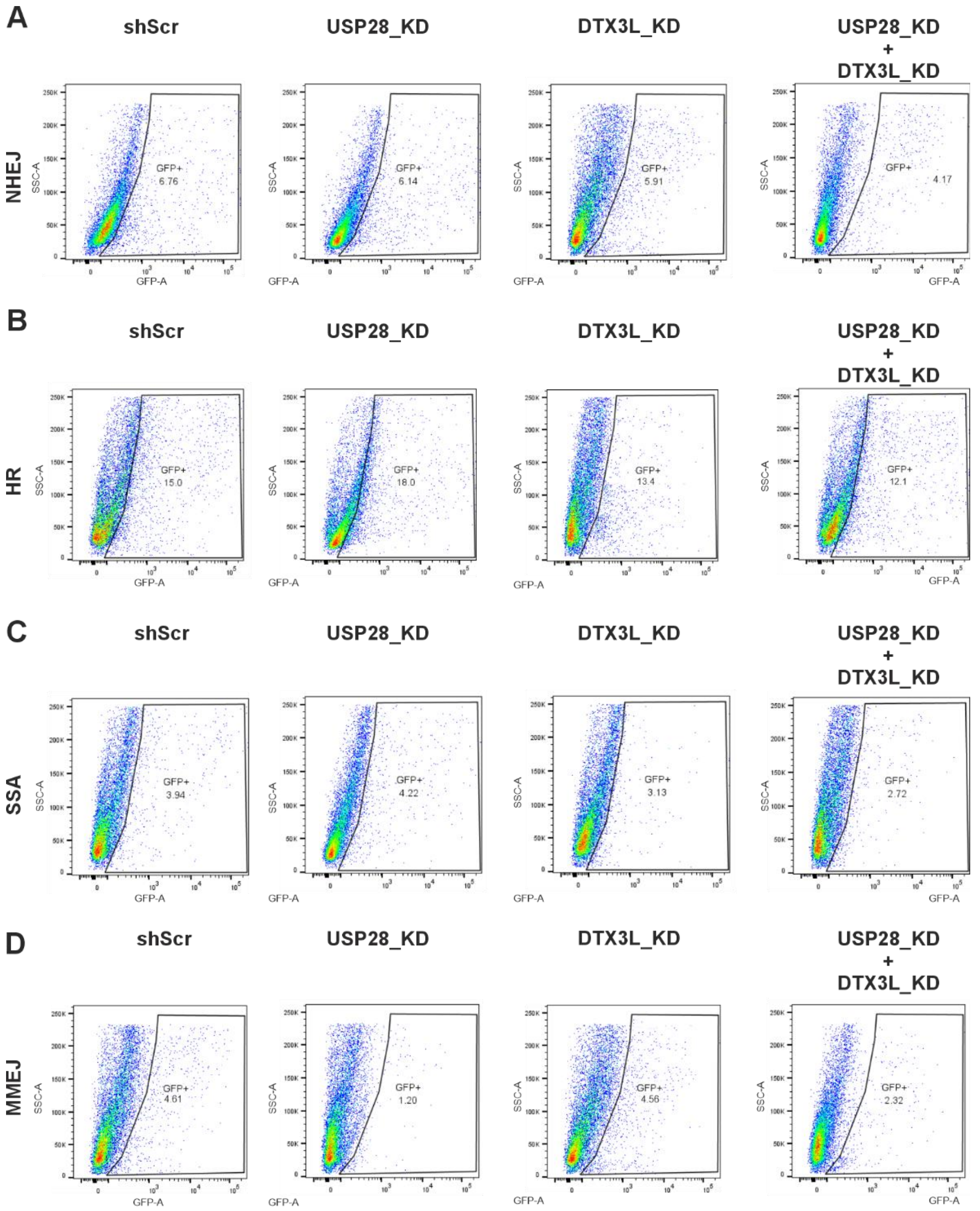


**Figure S9. Representative flow cytometry images from DSB repair analysis**. Pathway-specific repair activities were measured in cells using EGFP reporter assay. MDA-MB-231 Scr, and USP28 KD cells transfected with shRNAs against DTX3L or empty vector were cultured for 24 h prior to nucleofection with a DNA mixture consisting of I-*Sce*I meganuclease expression plasmid pCMV-I-Sce-I, balancing plasmid pBS (determination of repair frequency) or wild-type EGFP expression plasmid (determination of transfection efficiency), and DSB repair substrate to evaluate NHEJ **(A)**, HR **(B)**, SSA **(C)**, and MMEJ **(D)**

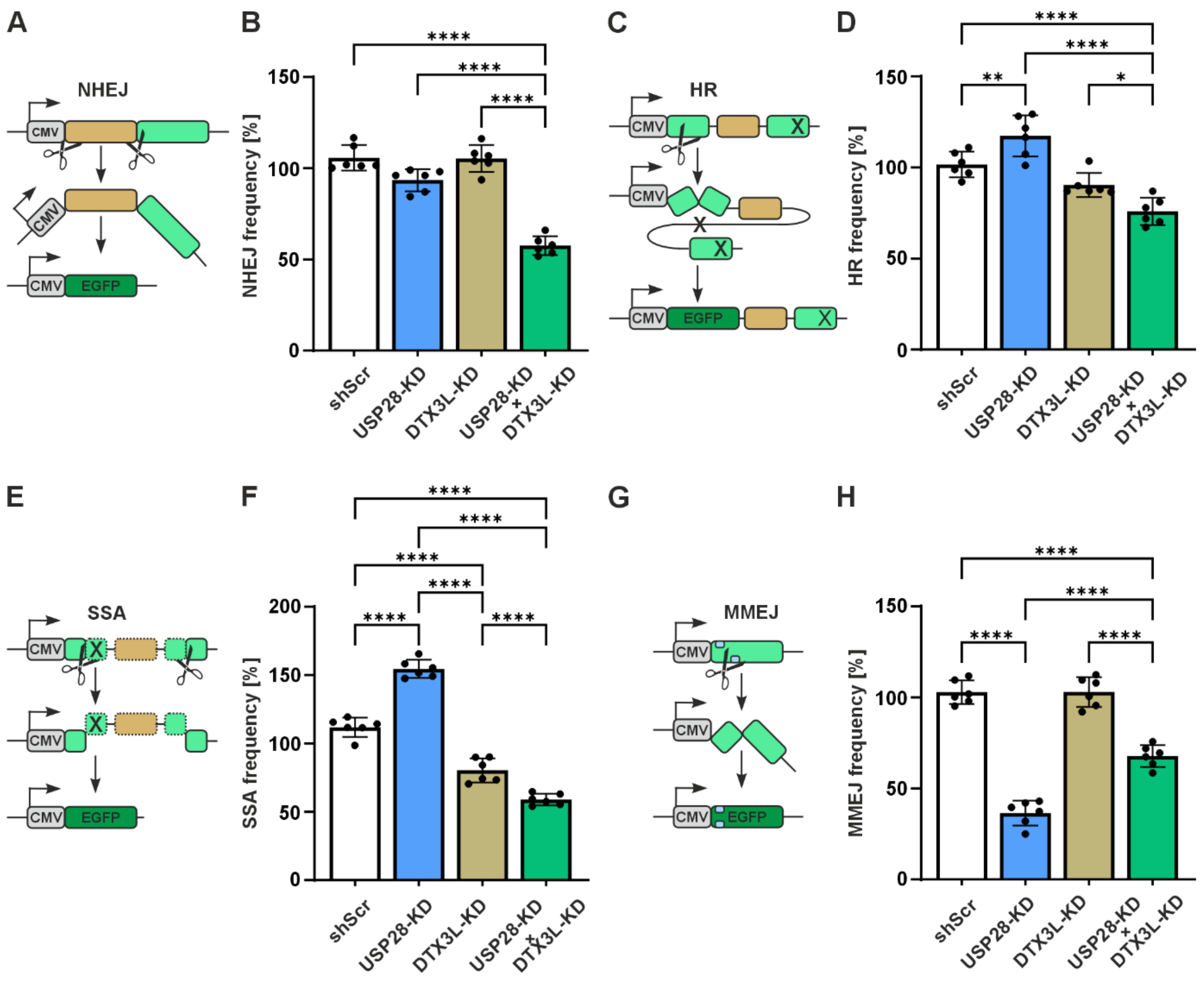


**Figure S10.** DSB repair analysis. Pathway-specific repair activities were measured in cells using EGFP reporter assay. MDA-MB237 Scr, and USP28 KD1 cells transfected with shRNAs against DTX3L or empty vector were cultured for 24 h prior to nucleofection with a DNA mixture consisting of I-*Sce*I meganuclease expression plasmid pCMV-I-Sce-I, balancing plasmid pBS (determination of repair frequency) or wild-type EGFP expression plasmid (determination of transfection efficiency), and DSB repair substrate to evaluate NHEJ **(A,B)**, HR **(C,D)**, SSA **(E,F)** or MMEJ **(G, H)**. Percentages of EGFP-positive cells were measured 48 h later and normalized to the individually determined transfection efficiencies to allow calculation of DSB repair frequencies. Mean values for the MDA-MB237 Scr cells (shScr) were defined as 100%. Data are mean +/- SD from 4-6 measurements. The statistical significance of differences was determined using ordinary one-way ANOVA. *$p<0.05$, **$p<0.01$ ***$p<0.001$.

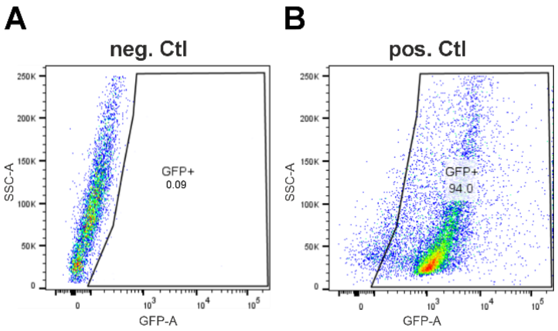


**Figure S11.** Controls for flow cytometry. MDA-MB-231 cells were transfected either with empty vector control (A) or wild-type EGFP expression plasmid (B). After transfection, EGFP-positive cell fractions were quantified by flow cytometry.

**Table S1. List of antibodies used in the study.**

| Primary Antibodies | Origin | Clonality | Dilution factor | Company | Catalog nr. |
|---|---|---|---|---|---|
| Ubiquitin | Mouse | monoclonal | 1:1.000 | BioLegend | #646304 |
| HIF-1α | Mouse | monoclonal | 1:1.000 | BD Bioscience | #610959 |
| p53 | Mouse | monoclonal | 1:1.000 | CST | #48818 |
| HDAC1 | Mouse | monoclonal | 1:1.000 | CST | #5356 |
| DTX3L | Mouse | monoclonal | 1:500 (WB)<br>1:100 (IF) | Santa Cruz | #sc-514776 |
| HA-Tag | Mouse | monoclonal | 1:1.000 | Santa Cruz | #sc7392 |
| FlagM2 | Mouse | monoclonal | 1:5.000 | Sigma-Aldrich | #F1804 |
| α-tubulin | Mouse | monoclonal | 1:10.000 | Sigma-Aldrich | #T5168 |
| USP28 | Rabbit | polyclonal | 1:1.000(WB)<br>1:100 (IF) | Sigma-Aldrich | #HPA006778 |
| DTX3L | Rabbit | polyclonal | 1:1.000 (WB) | Sigma-Aldrich | #HPA010570 |
| 53BP1 | Rabbit | polyclonal | 1:1.000 | Novus Biologicals | #NB100-304 |
| c-MYC | Rabbit | polyclonal | 1:500 | Santa Cruz | #sc-788 |
| **Secondary Antibodies** | **Origin** | | **Dilution factor** | **Company** | **Catalog nr.** |
| Mouse-HRP | Goat | | 1:5.000 | Bio-Rad | #1706516 |
| Rabbit-HRP | Goat | | 1:5.000 | Bio-Rad | #1706515 |
| Mouse-HRP | Goat | | 1:1.000 | DAKO | #P044701-2 |
| Alexa-Fluor 488 Mouse | Goat | | 1:1.000 (IF) | Invitrogen | #A-11001 |
| Alexa-Fluor 546 Rabbit | Goat | | 1:1.000 (IF) | Invitrogen | #A-11035 |

**Table S2. Primers used in the qRT-PCR analyses.**

| Gene | Forward Primer (5′→ 3′) | Reverse Primer (5′→ 3′) |
|---|---|---|
| *DTX3L* | CCAGGTTATGAGTCCTTTGGCAC | TGCAGTTCGCTGTATTCCAGGG |
| *USP28* | TGAAGCTCTGAAGGCCAGTAATG GTGAC | CTCCAGTAGACTCAAAGCAATG GCAGCCT |
| *ACTB* | GTTGTCGACGACGAGCG | GCACAGAGCCTCGCCTT |
